# Tracking eco-evolutionary dynamics among lung pathobiome members from children with cystic fibrosis

**DOI:** 10.64898/2026.01.29.702708

**Authors:** Lukas Schwyter, Roger Kuratli, Serena Provveduto, Phoebe Do Carmo Silva, Philipp Latzin, Freya Harrison, Markus Hilty, Rolf Kümmerli

## Abstract

Polymicrobial lung infections are common in individuals with cystic fibrosis (CF). Pathogen communities typically follow an ecological succession in which early colonizers such as *Haemophilus influenzae* and *Staphylococcus aureus* are later joined by *Pseudomonas aeruginosa*. Although adaptation to the lung environment is well described for single pathogens in adult people with CF, much less is known about the role of pathogen interactions and how changes in individual pathogens influence community dynamics at the early stages of disease. To address these questions, we combined genome sequencing with phenotypic screens and pathogen interaction assays using longitudinal clinical isolates (19 *P. aeruginosa*, 44 *S. aureus*, and 21 *H. influenzae*) collected from 23 children with CF (0–7.9 years of age) enrolled in the SCILD (Swiss CF Infant Lung Development) cohort. Our analysis revealed that early pathogen communities are characterized by a combination of strain turnover and persistence of isolates undergoing first steps of within-host evolution. Most notably, quorum-sensing–deficient *P. aeruginosa* variants repeatedly emerged, showing reduced protease production and diminished inhibition of *S. aureus* and *H. influenzae*. These changes indicate that *P. aeruginosa* becomes less antagonistic towards co-occurring pathogens, possibly promoting community stability. Together, our results show that ecological and evolutionary dynamics between pathobiome members may play an underappreciated role in shaping CF lung disease during early childhood.

## Introduction

The lungs of people with cystic fibrosis (CF) are colonized by a diverse consortium of pathogenic bacteria. Lung colonization starts at a very young age and is reminiscent of an ecological succession, whereby early colonizers such as *Haemophilus influenzae* and *Staphylococcus aureus* are followed by late colonizers such as *Pseudomonas aeruginosa* and species of the *Burkholderia cepacia* complex ^1,2^. Importantly, early-arriving species from the oropharyngeal flora may modify the airway microenvironment (e.g., oxygen concentration, pH, and nutrient availability), thereby constructing a niche that facilitates subsequent pathogen establishment and persistence ^3^. The pathobiome is further shaped by a range of other microbes including *Stenotrophomonas maltophila*, *Achromobacter xylosoxidans*, and non-tuberculous *Mycobacteria* species as well as fungal species of the genus *Aspergillus* ^4,5^. Although pathobiome composition differs across people with CF, *H. influenzae*, *S. aureus* and *P. aeruginosa* often co-occur, potentially fostering within-host species interactions and evolution^6^.

While certain pathogens are relatively benign others like *P. aeruginosa* and *B. cepacia* complex species are strongly associated with declining health and are often responsible for respiratory deterioration and death at the final stage of the disease ^7,8^. Because of these detrimental effects, there has been great interest in understanding the evolution of these pathogens and their interactions with the host. *P. aeruginosa* has become the major focus of evolutionary research because of its high prevalence (up to 80%) among adult people with CF ^9^. Key insights from this body of research revealed that people with CF are typically colonized by a single strain, which undergoes diversification and adaptive evolution in the lungs over decades ^10–12^. Common adaptation within the lung include the development of slow-growing phenotypes associated with increased mucoidy, biofilm formation and antibiotic resistance ^13–16^. Furthermore, a reduction in acute virulence factor production due to mutations in quorum sensing and other virulence factor regulators are also commonly observed ^17–20^.

Some of the evolutionary patterns, particularly the loss of virulence factor production, have led to a debate about whether *P. aeruginosa* evolution predominantly reflects adaptation to the host environment or whether it is driven by competition among diverging bacterial lineages within the host ^21^. While the most likely answer is that both factors influence evolutionary trajectories of *P. aeruginosa*, the debate neglects the possibility that *P. aeruginosa* may also adapt to other members of the pathobiome ^5,22^. Indeed, the other two prevalent pathogens, *S. aureus* and *H. influenzae*, are quite often considered bystanders ^23^ in CF lung infections and their interactions with *P. aeruginosa* lung isolates are poorly understood. Although *S. aureus* and *H. influenzae* may have less of an impact on disease progression, they can still significantly influence the lung environment and the ecological dynamics between pathogens therein ^24,25^. Changes in interaction patterns between pathogens could be due to direct or indirect effects. Indirect effects could arise when each pathogen independently adapts to the lung environment with the corresponding adaptations impacting ecological dynamics between species. Conversely, direct effects could arise when pathogens interact with one another through cell-to-cell contact or secreted compounds and specifically adapt to the presence of other community members ^26^.

In this study, we investigate the interaction patterns among *P. aeruginosa*, *S. aureus* and *H. influenzae* strains isolated from CF children of the SCILD (Swiss Cystic Fibrosis Infant Lung Development) cohort ^27^. We aimed to examine whether phenotypic changes in one species can affect the interaction dynamics with the other members of the pathobiome. We first sequenced the genomes of the collected isolates to capture phylogenetic associations and genetic diversity within each pathogen species. We then screened all isolates for a range of virulence factor phenotypes including protease production, siderophore production, biofilm formation, mucoidy, and hemolysis, and scored their antibiotic resistance profiles. These analyses allowed us to track genetic and phenotypic changes in all three pathogens across age in children with CF. Subsequently, we conducted a large-scale (spent) supernatant assay to quantify interactions patterns via secreted exoproducts between the three pathogens across sampling time windows. Finally, we combined the genetic, phenotypic and interaction data to derive mechanistic and fitness-based models of how change within one of the CF pathogens may affect the ecological dynamics with the other members of the pathobiome.

## Results

### Longitudinal sampling of *P. aeruginosa, S. aureus*, and *H. influenzae* isolates across 23 children with cystic fibrosis

From the SCILD cohort, we obtained 19 *P. aeruginosa*, 44 *S. aureus*, and 21 *H. influenzae* isolates retrieved from oropharyngeal swaps collected from 23 children with CF (aged between 0 and 7.9 years, Figure 1, Table S1) ^27^. All children underwent regular clinical evaluations, during which oropharyngeal swabs were obtained to monitor airway colonization. Based on the isolate sampling, *S. aureus* was the most prevalent pathogen present in 65% of all children with CF, followed by *H. influenzae* (57 %) and *P. aeruginosa* (43%). The selection of isolates, therefore, roughly represents the well-known assembly of a multi-species pathobiome in the CF airways.

**Figure 1.**
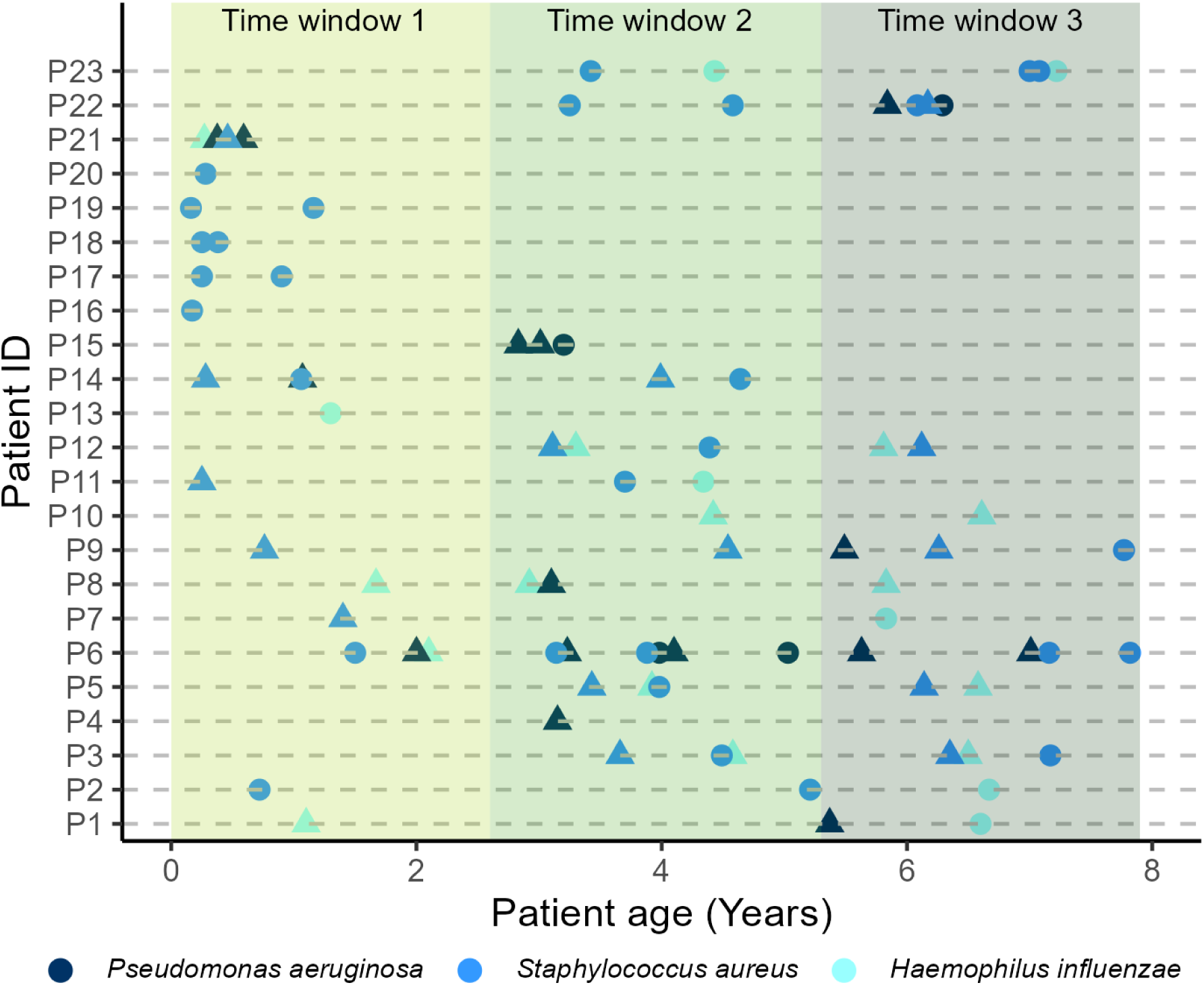
Distribution of clinical bacterial isolates collected from 23 children with cystic fibrosis. Scatter plot showing the distribution of pathogen isolates across children with CF and their age, separated by pathogen species (color). Isolates were allocated to time windows (early, intermediate, late isolates), indicated by the different shadings. For all 84 isolates, genomes were sequenced, and virulence and antibiotic resistance phenotypes were assessed. A subset of randomly selected isolates (triangles) was used to establish interaction networks across species and time windows.

Since pathogen prevalence and sampling scheme were heterogenous across children and time, the isolate collection does not permit us to directly study pathogen interactions and potential (evolutionary) changes within children. However, under the assumption that changes in the lung environment and thereby selection pressures are similar across people with CF ^28^, we pooled isolates across children and divided them into three discrete time windows, separating samples into early (time window T1: 0–2.6 years; isolates: 25), intermediate (time window T2: 2.6–5.3 years; isolates: 30), and late (time window T3: 5.3–7.9 years; isolates: 29) isolates (Figure 1). In a first step, we sequenced the genomes and collected data on virulence factor phenotypes and antibiotic resistance profiles for all isolates to capture genetic and phenotypic diversity among isolates for each pathogen separately.

### *P. aeruginosa* genomes increase in size with age and cluster by serogroup and child

We examined the phylogenetic relationship among the 19 clinical *P. aeruginosa* isolates based on 5111 shared core genes and included two well-characterized laboratory strains (PAO1, PA14) as references (Figure 2A). We conducted an association index analysis to test whether isolates cluster by serogroup, time window, and individual (minimum requirement: N = 3 isolates per category). We observed four different serogroups, whereby O6 was most frequent (11/19) followed by O3 (3/19), O9 (3/19) and O11 (2/19). The serotypes O3 (p = 0.0008), O6 (p = 0.0010) and O9 (p = 0.0009) all showed significant clustering, highlighting that serotype is a reliable phylogenetic marker. Conversely, there was little clustering by time window, with significant clustering only arising in T1 (p = 0.0114), but not in T2 (p = 0.0701) and T3 (p =0.0827), whereby the clustering for T1 is probably a confounding effect because two out of the four clones were isolated from the same child. Overall, the low clustering by time window is expected as each child likely picked up a distinct clone. In support of this, we found significant clustering for the individuals P6 (p = 0.0096, N=7) and P15 (p = 0.0015, N=3) indicating the persistence of specific clones within individuals already at a young age.

**Figure 2.**
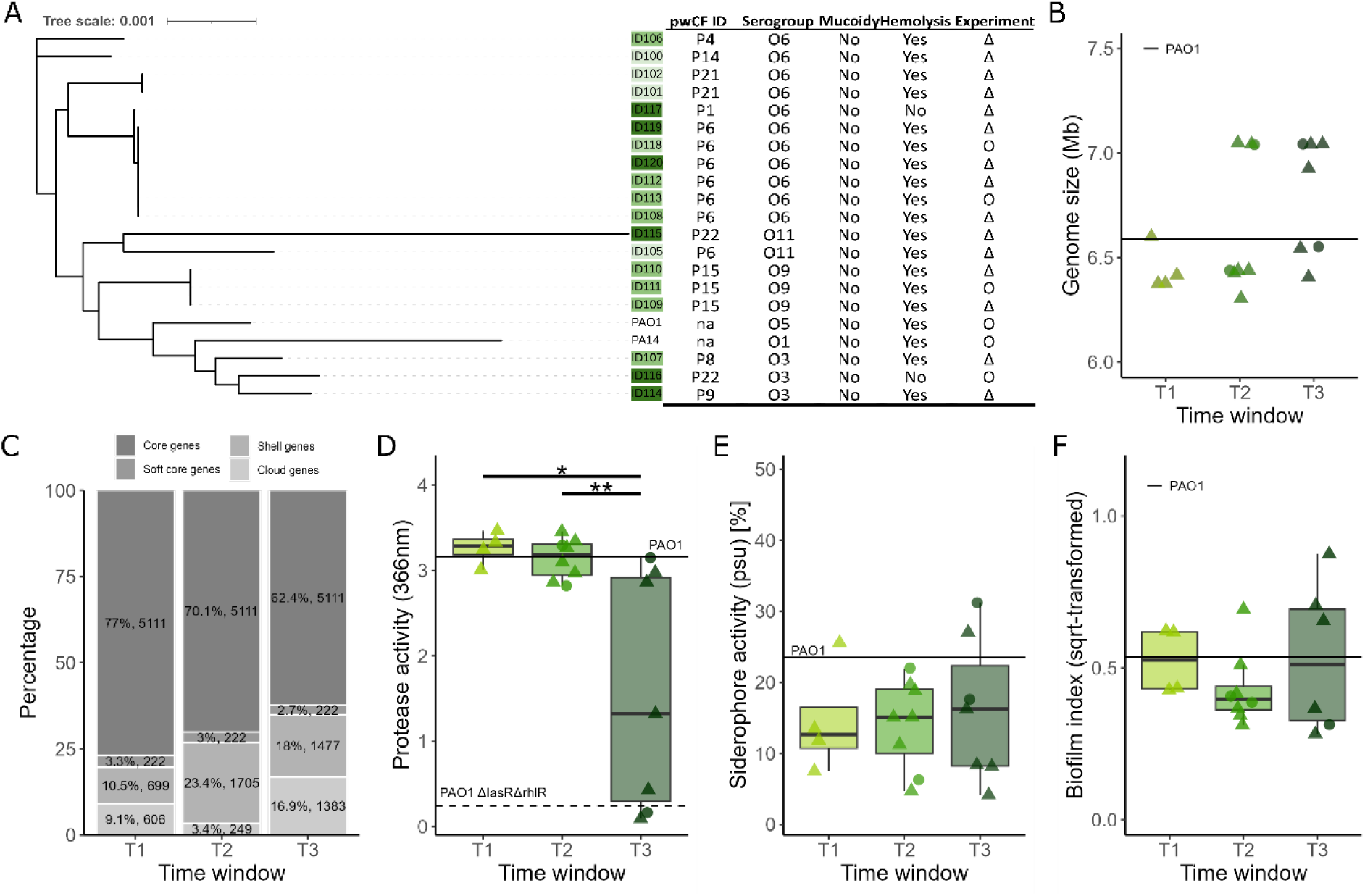
Phylogenetic relationship, genome composition and phenotypes of 19 *P. aeruginosa* isolates from children with CF. **(A)** Phylogenetic tree based on 5111 core genes of 19 clinical isolates and two well-studied *P. aeruginosa* lab strains (PAO1 and PA14). The table displays the ID of people with CF (pwCF), serogroup, as well as the presence/absence of mucoid and hemolytic phenotypes. **(B)** Genome size of clinical isolates in base pairs (bp) split by time window. **(C)** Average genome compositions of clinical isolates across time windows, with genes classified as core (present in 100% of isolates), soft-core (90–99%), shell (15–89%), and cloud (0–14%). **(D)** Protease activity of clinical isolates measured with the azocasein assay. **(E)** Siderophore production of the clinical isolates measured with the chrome azurol S (CAS) assay in ssBHI medium supplemented with 600 μM 2,2’-bipyridine. **(F)** Biofilm formation under static growth conditions measured with the crystal violate assay. Asterisks denote significance levels: ***P < 0.001; **P < 0.01; *P < 0.05.

When focusing on genome composition, we observed an increase in genome size over time (T1: 6.44 Mbp, T2: 6.65 Mbp and T3: 6.80 Mbp), with increasing fractions of cloud and shell genes being present in T2 and T3 isolates (Figure 2B). Specifically for child P6, who had strains present across T1, T2, and T3, we noted that the genome size increased from T1 (6.42 Mbp) to T2 and T3 (both 7.05 Mbp). To further explore genomic variation over time, we calculated Jaccard distance matrixes based on gene presence/absence across isolates. The analysis revealed increasing Jaccard distances, raising from 0.106 (T1) to 0.131 (T2) and 0.171 (T3), indicating progressive temporal genomic diversification among isolates from the same time window (Figure S1A).

To investigate functional aspects of this increase, we grouped all genes according to their COG groups ^29,30^ and counted the number of unique genes per time window (Figure S2A). We observed significant temporal shifts in functional gene content between T1 and T3. Notably, two COG categories showed significant decreases in the number of unique genes: category F (RNA processing and modification: mean change = −1.3; Dunn’s test: Z =-3.04, p = 0.0036) and J (translation, ribosomal structure and biogenesis: mean change= −3.4; Z =-3.21, p = 0.0020). In contrast, the largest increases were observed in category L (replication, recombination and repair: mean change = +16; Z = 5.2, p < 0.0001), followed by N (cell motility: mean change= +9.7; Z = 4.4, p = 0.0003) and U (intracellular trafficking, secretion, and vesicular transport: mean change = +8.1; Z = 4.78, p < 0.0001). These findings point towards systematic changes in PA genome composition with increasing child age.

### *P. aeruginosa* isolates from the last time window show reduced protease production

We assessed five phenotypic traits involved with virulence (proteases, siderophores, biofilm, mucoidy and hemolysis) when *P. aeruginosa* isolates were grown in a medium permitting growth of all three species of interest (BHI broth supplemented with NAD and hemin: ssBHI, with the siderophore assay done in iron depleted media: ssBHI + 600 µM 2,2′-bipyridine) and tested whether these traits varied across isolates and time windows. We observed a significant reduction in protease activity across time windows (linear mixed-effects model: F_2, 16_ = 7.56, p = 0.0049, Figure 2D), with isolates from T3 producing on average significantly less proteases compared to isolates from T1 and T2 (pairwise comparisons, T1 vs. T2: estimate = 0.122 ± 0.534, t_16_ = 0.23, p = 0.9716; T1 vs. T3: estimate = 1.691 ± 0.547, t_16_ = 3.09, p = 0.0181; T2 vs. T3: estimate = 1.569 ± 0.451, t_16_ = 3.48, p = 0.0083). In contrast, we did not find significant differences in average siderophore iron chelating activity (Figure 2E; linear mixed-effects model: F_2, 16_ = 0.12, p = 0.8915) and biofilm formation (Figure 2F; *F*_2, 15_ = 0.84, p = 0.4510) across time windows. Instead, we observed increased variation in siderophore production among isolates at T3 with some of the isolates producing very low amounts. When comparing phenotypes across serogroups (O3, O6, O9, O11), only biofilm formation differed significantly (Figure S3; *F*_3,14_ = 6.87, p = 0.0045). However, these serogroup analyses had low statistical power due to small sample sizes. None of the isolates had a mucoid phenotype when grown on ssBHI agar and 17 out of 19 isolates were hemolysis positive when grown on Columbia agar supplemented with sheep blood (Figure 2A). Altogether, our phenotypic screen revealed overall lower protease production levels and increased variation in siderophore production levels among isolates from older children (T3).

### *S. aureus* genomes cluster by *spa* type and by child

We explored the phylogenetic relationship among the 44 clinical *S. aureus* isolates and four laboratory reference strains (Newman, JE2, MW2, and 6850) based on 1947 core genes (Figure 3A). Our analysis revealed 22 different *spa* types (staphylococcal protein A, an important virulence factor involved in human immune evasion), indicating that children with CF are colonized by a diverse set of *S. aureus* strains ^31^. The association index analysis showed significant clustering for all *spa* types with N ≥ 3: t2 (p = 0.0048), t267 (p = 0.0045), t349 (p = 0.0027), t688 (p = 0.0049), and t84 (p = 0.0072). We further found significant phylogenetic clustering of isolates from time window for T1 (p = 0.018) and T2 (p = 0.0165), but not T3 (p = 0.2408), a finding that could again be confounded by the repeated sampling of genetically similar isolates from the same child (Figure 1). Indeed, we found significant phylogenetic clustering by child in four cases (P5: p = 0.0034, N=3; P6: p = 0.0054, N=5; P9: p = 0.0077, N=4; P14: p = 0.0052, N=4, with all these individuals maintaining consistent *spa* types across time. In the remaining three cases, no evidence for within-individual clustering occurred (P3: p = 0.6588, N = 4; P9: p = 0.8331, N = 4; P12: p = 0.3262, N = 3) consisting with the observation that *spa* types changed over time. Taken together, these results confirm *spa* type as a phylogenetic marker and suggest that both retention of the same clone and strain turnover can occur depending on the child ^32,33^.

**Figure 3.**
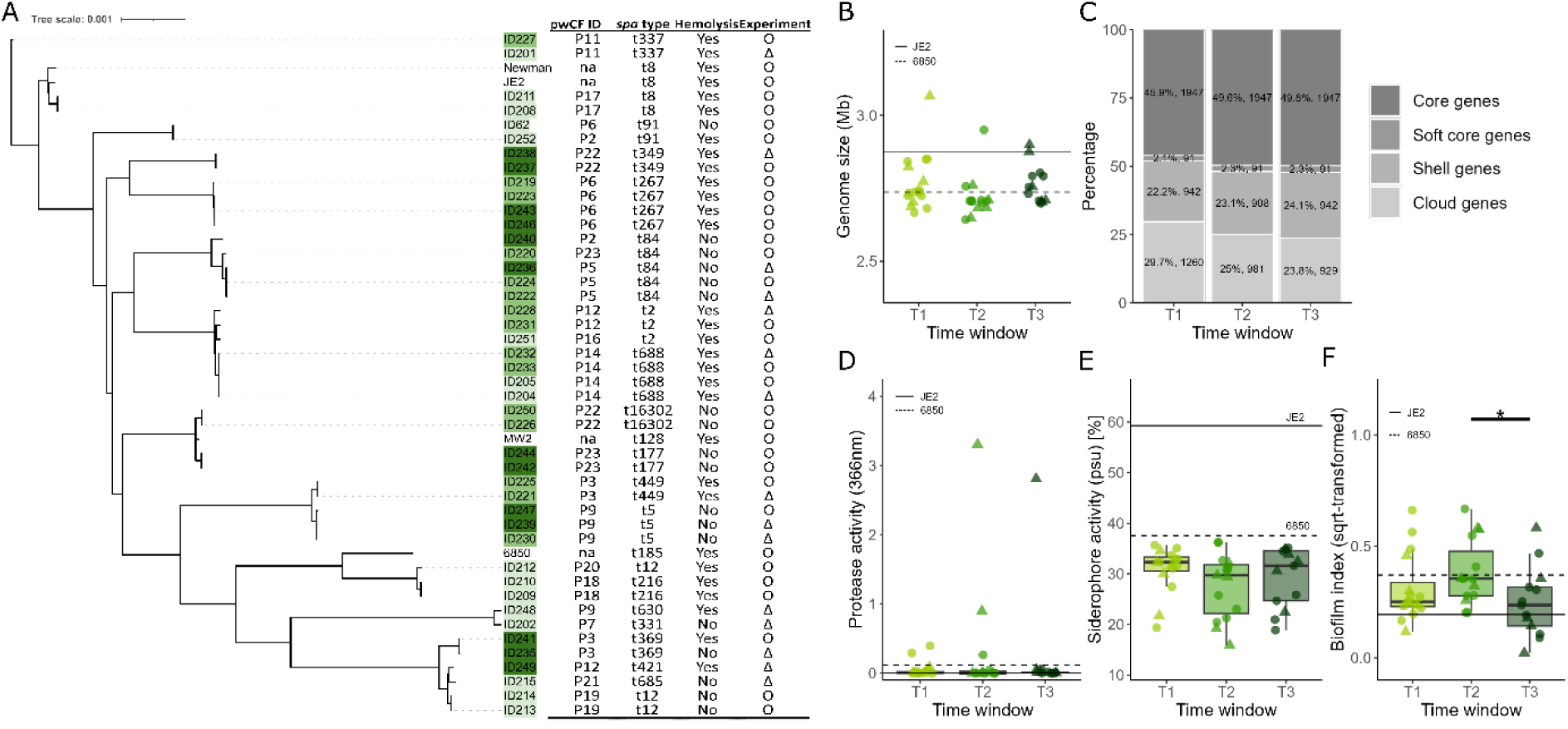
Phylogenetic relationship, genome composition and phenotypes of 44 *S. aureus* isolates from children with CF. **(A)** Phylogenetic tree based on 1947 core genes of clinical S*. aureus* isolates and four well-studied *S. aureus* laboratory strains (Newman, JE2, MW2, and 6850). The table displays the ID of people with CF (pwCF), *spa* type, and hemolytic activity of each isolate. **(B)** Genome size (in base pairs) of the clinical isolates, grouped by time window. **(C)** Average genome composition of clinical isolates, with genes classified as core (present in 100% of isolates), soft-core (90–99%), shell (15–89%), and cloud (0–14%). **(D)** Protease activity of clinical isolates measured using the azocasein assay. **(E)** Siderophore production of clinical isolates measured using the chrome azurol S (CAS) assay in ssBHI medium supplemented with 75 μM 2,2’-bipyridine. **(F)** Biofilm formation under static growth conditions measured using the crystal violet assay. Asterisks denote significance levels: ***P < 0.001; **P < 0.01; *P < 0.05.

We found no significant differences in genome size and composition across time windows (Kruskal–Wallis test: χ² = 5.15, *df* = 2, p = 0.0762). Average genome size was 2.75 Mbp and varied between 2.66 Mbp and 2.89 Mbp (Figures 3B+C). Consistent with this, Jaccard distances remained relatively constant over time (T1: 0.228; T2: 0.187; T3: 0.223, Figure S1C).

Although overall genome features (i.e., frequency of core, soft, shell, cloud genes) did not change over time, we observed significant temporal variation in specific functional COG categories (Figure S2B). The most prominent reductions in unique gene counts occurred in category T (signal transduction mechanisms, which includes loci encoding the *agr* quorum sensing system: mean change = −8.1; Dunn’s test: Z =-4.99, p < 0.0001), category H (coenzyme transport and metabolism: mean change = −5; Z =-2.62, p = 0.0086) and category G (carbohydrate transport and metabolism: mean change = −4.7; Z =-4.10, p = 0.0007), whereas the highest increase in unique gene counts occurred in category K (transcription mean change = 2.40; Z = 4.66, *p* < 0.0001). These findings suggest that SA genome size remains stable over time, but shifts in the abundance of specific gene categories correlate with child age.

### *S. aureus* isolates show variation in siderophore and biofilm formation across time

For *S. aureus* isolates, we assessed the phenotypes for proteases, siderophores, biofilm, and hemolysis, and further examined the occurrence (qualitative presence/absence) of small colony variants (SCV; presence of colonies with a diameter <50% that of the typical *S. aureus* diameter). These assays revealed that almost all *S. aureus* isolates lacked detectable protease activity under our assay conditions (Figure 3D). For both siderophore production and biofilm formation, we observed substantial phenotypic variation across isolates within each time window. There were no significant differences in siderophore production between time windows and no evidence of a temporal trend (linear mixed-effects model: F_2, 39.99_ = 1.17, p = 0.3214, Figure 3C). In contrast, biofilm formation differed significantly across time windows (F_2, 41.05_ = 3.98, p = 0.0263) with biofilm formation being higher at time window T2 compared to T3 (estimate = 0.147 ± 0.053, t_40.9_ = 2.75, p = 0.0238, Figure 3D). Regarding hemolysis, we observed hemolysis positive and negative isolates in all time windows (T1: 11 positive versus 5 negative isolates; T2: 9 versus 6; T3: 7 versus 6), indicating persisting heterogeneity in this trait over time. Streaking the isolates on mannitol-salt agar revealed the presence of SCVs in almost all isolates (Figure S4). Altogether, these analyses reveal high phenotypic diversity among SA isolates from pediatric CF patching, matching the high genetic diversity observed (Figure 3A).

### *H. influenzae* isolates show no specific phylogenetic clustering yet exhibit a temporal accumulation of shell and cloud genes

We examined the phylogenetic relationship among the 20 sequenced clinical *H. influenzae* isolates (one isolate was lost due to poor sequencing quality) and three laboratory strains (ATCC 49247, ATCC 10211, and 86-028NP) based on 1196 core genes (Figure 4A). The association index revealed neither a significant clustering according to time window (T1: p = 0.2539; T2: p = 0.1510; T3: p = 0.1826) nor according to individual (P8: p = 0.5395, N = 3). These results suggest that children with CF are colonized by a diverse range of *H. influenzae* isolates with substantial within-individual turnover, consistent with prior observations of frequent strain replacement and only rare prolonged carriage in adult persons with CF ^34^.

**Figure 4.**
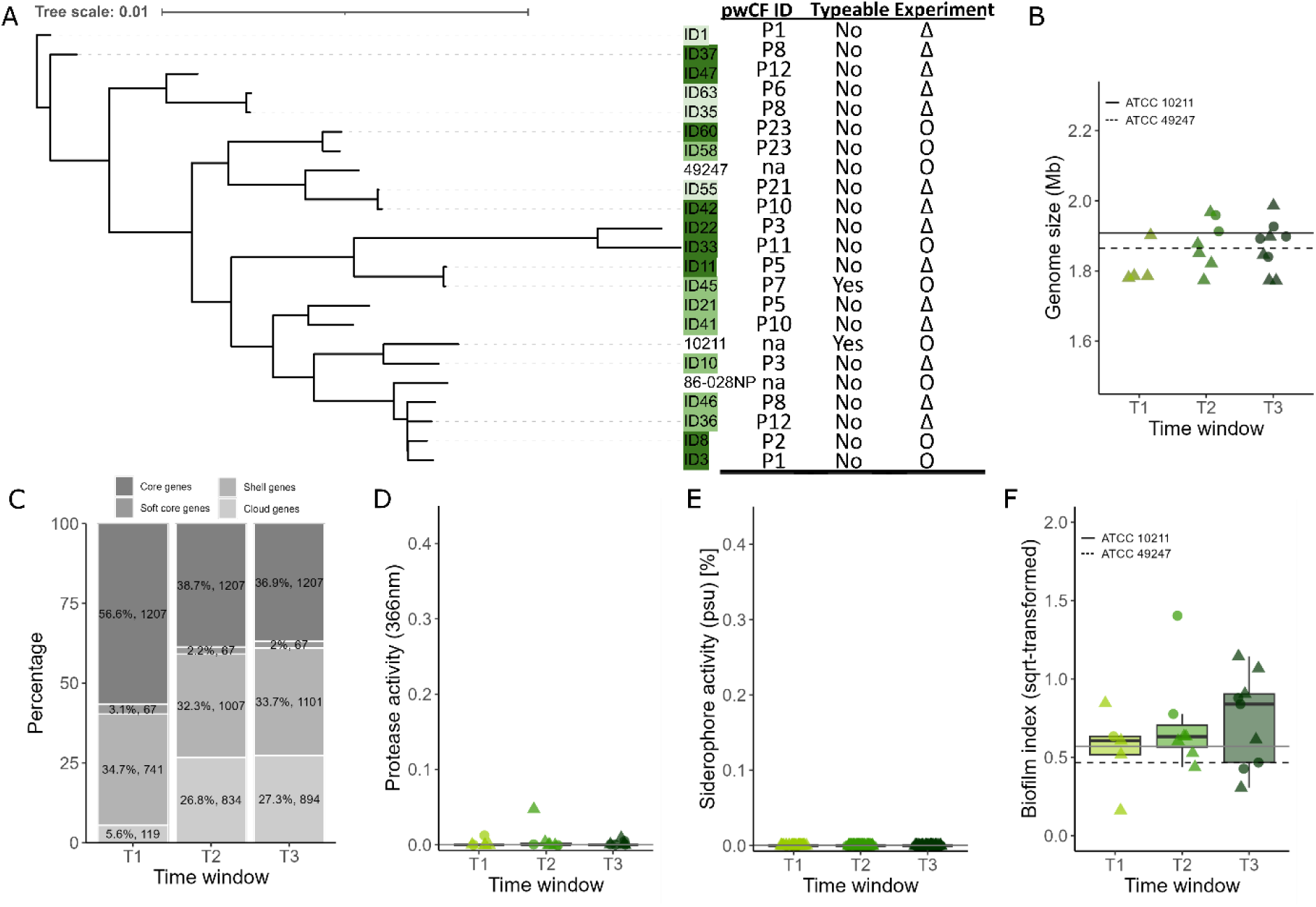
Phylogenetic relationship, genome composition and phenotypes of 21 *H. influenzae* isolates from children with CF. **(A)** Phylogenetic tree based on 1196 core genes of clinical *H. influenzae* isolates and three well-studied laboratory strains (86-028NP, ATCC 10211, and ATCC 49247). The table displays the ID of people with CF (pwCF) and the capsule presence (typeable vs non-typeable). Note that one isolate had to be excluded from genetic analysis due to poor sequence qualities. **(B)** Genome size (in base pairs) of the clinical isolates, grouped by time window. **(C)** Average genome composition of clinical isolates, with genes classified as core (present in 100% of isolates), soft-core (90–99%), shell (15–89%), and cloud (0–14%). **(D)** Protease activity of clinical isolates measured using the azocasein assay. **(E)** Siderophore production of clinical isolates measured using the chrome azurol S (CAS) assay in ssBHI medium supplemented with 2,2’-bipyridine to a final concentration of 75 μM. **(F)** Biofilm formation under static growth conditions measured using the crystal violet assay. Asterisks denote significance levels: ***P < 0.001; **P < 0.01; *P < 0.05.

Comparative genomic analysis revealed stable genome sizes across the time windows (Kruskal–Wallis test: χ² = 1.49, df = 2, p = 0.4758, T1: 1.82 Mbp; T2: 1.88 Mbp; T3: 1.87 Mbp, Figure 4B). In contrast, we detected substantial changes in genome composition (Figure 4C). Particularly, there were temporal increases in both “shell” (T1: 741; T2: 1007; T3: 1101) and “cloud” (T1: 119; T2: 834; T3: 894) genes. These shifts were mirrored in the Jaccard distances (Figure S1C) that increased from T1 (0.190) to T2 (0.300) and T3 (0.314). Similarly, the COG category analysis revealed increases in unique gene counts for various categories, whereas no decreases occurred. The largest increases were observed in category J (translation, ribosomal structure and biogenesis: mean change = 12.5, Dunn’s test: Z = 4.51, p < 0.0001), followed by category L (replication, recombination and repair: mean change = 10.9, Z = 3.37, p = 0.0011;), and E (amino acid transport and metabolism: mean change = 7.3, Z = 3.84, p = 0.0004). These analyses show that the *H. influenzae* genome composition differs fundamentally according to time window characterized by an enrichment in accessory genome elements over time.

### *H. influenzae* isolates are predominantly non-typeable and show low phenotypic variation

As for the other two species, we assessed the phenotypes for proteases, siderophores, and biofilm (but not hemolysis because *H. influenzae* is non-hemolytic). There were no significant changes in phenotypes across time windows. Specifically, most isolates did not produce any detectable level of proteases (Figure 4D) and none of the isolates produced siderophores (Figure 4E). Biofilm formation, however, was consistently detected in almost all isolates and biofilm formation remained stable across time windows (Figure 4F, linear mixed-model effect: F_2, 18.12_ = 0.82, p = 0.4556). We further assessed the typeability of isolates because the presence or absence of a capsule is an important determinant of virulence and immune recognition. While encapsulated strains are associated with invasive disease due to their ability to resist phagocytosis, non-typeable strains are better adapted for colonization of mucosal surfaces ^35,36^. Among the 21 isolates, only one was typeable, whereas the remaining 20 isolates were non-typeable. Overall, these data suggest that non-typeable *H. influenzae* isolates predominate in children with CF ^37^, which is not surprising given that children in Switzerland are routinely vaccinated against encapsulated *H. influenzae* type b. Isolates exhibited temporally consistent phenotypic characteristics

### Isolates vary considerably in their antibiotic resistance profiles

Children with CF undergo antibiotic treatment, particularly when infected with *P. aeruginosa*, but also against *S. aureus* and *H. influenzae* infection. Since antibiotic treatment can select for resistance in both targeted and non-targeted species ^38^, we assessed the susceptibility of all isolates towards a panel of six antibiotics (ampicillin, ciprofloxaxin, colistin, meropenem, piperacillin-tazobactam, tobramycin) in ssBHI (Table S2). Specifically, we measured the IC_90_ values (antibiotic concentration required to inhibit pathogen growth by 90%) and tested whether they change over time. Since our culturing conditions differ from standard antibiotic sensitivity tests, we compared IC90 values of our clinical isolates to those of reference strains for which antibiotic resistance profiles are determined according to EUCAST guidelines.

For *P. aeruginosa*, there were no significant temporal changes in IC_90_ values for any of the six antibiotics tested (Figure S3D, Figure S5A). All isolates showed high IC_90_ values for ampicillin similar to our PAO1 reference strain and consistent with the known intrinsic resistance of *P. aeruginosa* to this antibiotic due to the presence of endogenous beta-lactamase genes^39^. For the other five antibiotics, IC_90_ values of our CF isolates were either similar (ciprofloxacin, piperacillin–tazobactam, tobramycin) or below (colistin, meropenem) the IC_90_ values of PAO1, indicating overall antibiotic susceptibility.

For *S. aureus*, significant temporal changes occurred for two antibiotics (Figure S5B). Ciprofloxacin IC_90_ values significantly decreased over time (linear mixed-effects model: F_2,41_ = 3.23, p = 0.0498, T1 vs. T3: estimate = 0.699 ± 0.276, *t*₄₁ = 2.53, *p* = 0.0461). When comparing IC_90_ values to the reference strains JE2 (MRSA, ciprofloxacin resistant) and 6850 (MSSA, ciprofloxacin susceptible), our CF isolates grouped more closely with 6850 than with JE2, suggesting widespread susceptibility. In contrast, tobramycin IC_90_ values significantly increased over time (F_2,40.2_ = 4.50, p = 0.0173; T1 vs. T3: estimate = –0.866 ± 0.292, t_40.9_ = –2.97, p = 0.0151). Several isolates, especially from T3, exceeded IC_90_ values of both references suggesting reduced susceptibility and increased resistance. For ampicillin and piperacillin–tazobactam, we observed a bimodal distribution of IC_90_ values with one fraction of isolates clustering with or exceeding values for JE2 (indicating resistance), whereas the other fraction of isolates being closer to 6850 (indicating susceptibility). Finally, IC_90_ values for meropenem were low overall (except for three isolates) suggesting widespread susceptibility.

For *H. influenzae*, no significant temporal changes were detected for any antibiotic (Figure S5C). When comparing the IC_90_ values for ampicillin and piperacillin–tazobactam to the reference strains ATCC 49247 (β-lactamase positive ^40^) and ATCC 10211 (β-lactamase negative ^41^), we observed that isolates covered the entire IC_90_ space between the two references. This result indicates high heterogeneity in IC_90_ values among *H. influenzae* CF lung isolates, from susceptible to resistant. There was less variation in IC_90_ values for ciprofloxacin, colistin, meropenem and tobramycin, with IC_90_ values being generally low, suggesting susceptibility. Altogether, we observed high variation in antibiotic susceptibility across CF lung isolates for all three species but very few directional temporal changes in IC_90_ values. These findings provide limited evidence for the evolution of antibiotic resistance over the course of pediatric CF lung infections, yet underline the high diversity of isolates that colonize the lung of children with CF.

### Mapping within-species interactions patterns over time

In a next step, we assessed the interaction patterns between isolates from the same and different species using a supernatant assay. For this experiment, we focused on five random isolates per species and time window (T1 = 5, T2 = 5, T3 = 5), except for *P. aeruginosa* (T1 = 4, T2 = 6, T3 = 5) and exposed all isolates to a mix of 62% fresh medium and 38% spent supernatants of all other isolates in a full matrix comprising 2025 interactions. We quantified the effect of the supernatant on overall growth (AUC, area under the growth curve) and lag-phase relative to the control medium (62% fresh medium and 38% saline buffer). In this section, we focus on within-species interactions. Closely related isolates often have similar nutrient requirements. Nutritional overlap can foster competition mediated by secreted toxic compounds detectable in supernatants. Alternatively, nutritional overlap can also foster the cooperative scavenging of nutrients (e.g., through siderophore and enzyme secretion) and supernatants may thus contain growth stimulatory effects ^42^.

Among *P. aeruginosa* isolates, we found that supernatants had generally strong inhibitory effects on other *P. aeruginosa* isolates (Figure 5 + Figure S6A). The strength of inhibition significantly decreased and thus growth increased over time (AUC, linear mixed-effects model: F_(2, 77.0_ = 3.47, p = 0.0362). The effect was largest between T1 and T3 (estimate = 0.267 ± 0.109, t_77_ = –2.45, p = 0.0432), whereas differences between adjacent time windows were not significant (T1 vs. T2: estimate = – 0.048 ± 0.104, t_77_ = –0.46, p = 0.8914; T2 vs. T3 (estimate = –0.219 ± 0.104, t_77_ = –2.10, p = 0.0961).

**Figure 5.**
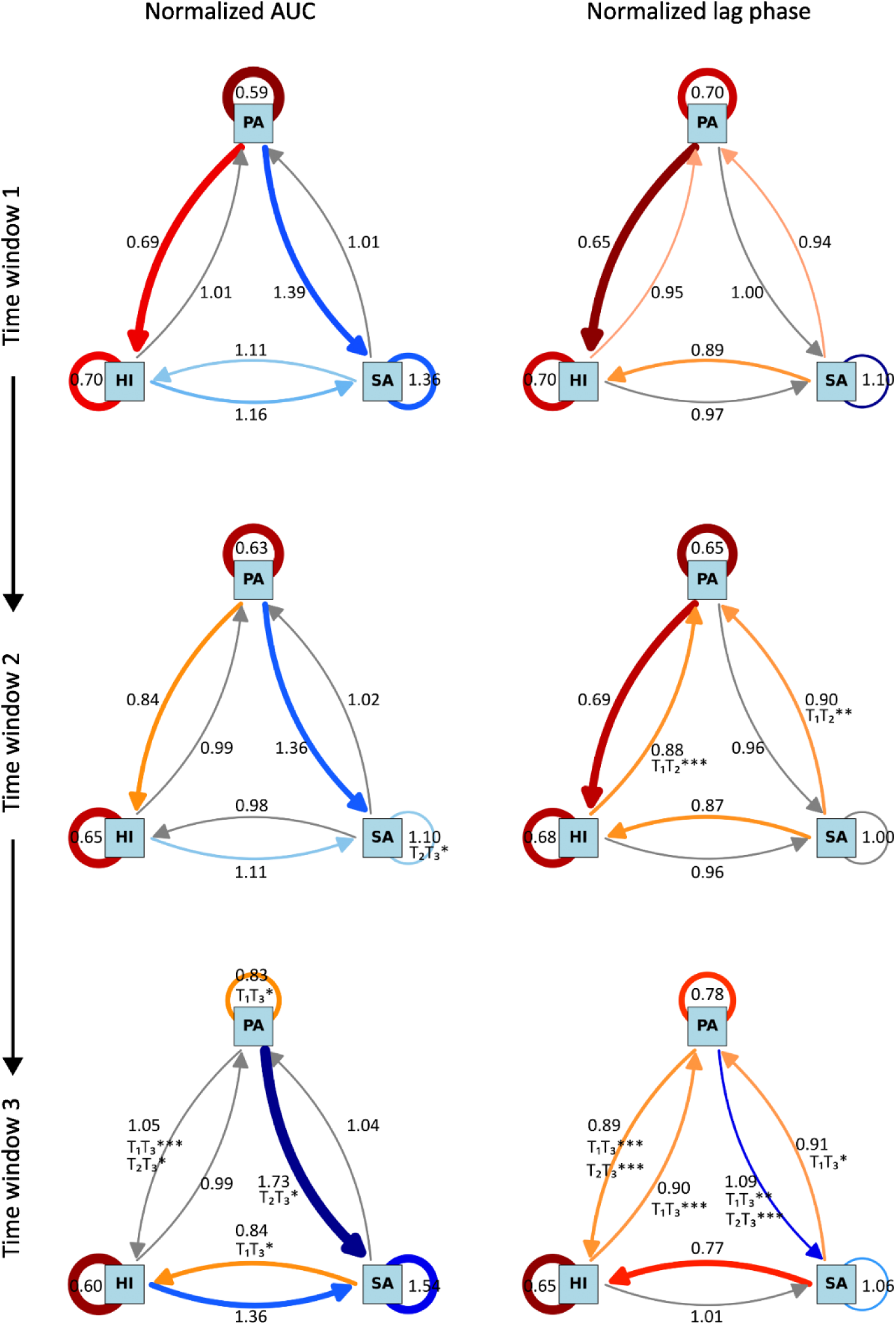
Significant changes in the interaction network between the three CF pathogens across time. **(A)** Interaction network at each time window between the three species using area under the curve (AUC) normalized to the control growth (63% ssBHI + 37% NaCl). The colors and thickness of the arrows indicate the strength of the effect: grey (1 ≤ AUC_norm_ ≤ 1.05) indicates no inhibition; blue (AUC_norm_ > 1.05) indicates growth promotion; orange (0.8 < AUC_norm_ < 1) indicates moderate inhibition; and red (AUC_norm_ < 0.8) indicates strong inhibition. The shading of the arrow colors represents the variance across samples with lighter colors indicating higher variance. **(B)** Interaction network at each time window between the three species using normalized lag phase as a proxy for short term interaction effects. The colors and thickness of the arrows indicate the strength of the effect: grey (1 ≤ Lag_norm_ ≤ 1.05) indicates no inhibition; blue (Lag_norm_ > 1.05) indicates growth promotion; orange (0.8 < Lag_norm_ < 1) indicates moderate inhibition; and red (Lag_norm_ < 0.8) indicates strong inhibition. The shading of the arrow colors represent the variance across the samples with lighter colors indicating higher variance. Asterisks denote significance levels: ***P < 0.001; **P < 0.01; *P < 0.05.

Among *S. aureus* isolates, we observed that supernatants had no inhibitory effect on other *S. aureus* isolates, but rather a positive effect on growth compared to the control condition (Figure 5 + Figure S6B), with the positive effects becoming slightly stronger over time (estimate = –0.447 ± 0.151, t_72.0_ = –2.97, p = 0.0112). Among *H. influenzae* isolates, supernatants had a consistently strong inhibitory effect on other *H. influenzae* isolates at all time windows (F_2, 69_ = 0.48, p = 0.6224, Figure S6C). Overall, there was high within-species variation in how supernatants of each of the three pathogens affected the growth of isolates of the same species (Figure S6). For *P. aeruginosa* and *H. influenzae,* within-species growth effects were often bimodal, meaning that certain supernatants were highly inhibitory whereas others had neutral or even positive growth effects on the receiving isolates. Taken together, our results highlight that isolates of the same pathogen species can negatively and positively interact with one another via secreted compounds, and that the strength of these interactions vary across time.

### Patterns of between-species pathogen interactions depend on species pairs and time

Next, we analyzed supernatant-based pairwise interactions between the three pathogen species.

(i) *P. aeruginosa* versus *H. influenzae* – *H. influenzae* typically colonizes the lungs first, while *P. aeruginosa* invades at later time windows. Given that *P. aeruginosa* is generally considered a competitive species, possessing an array of anticompetitor mechanisms ^43^ we expect this pathogen to be dominant over *H. influenzae*. Indeed, we observed that the supernatant of *P. aeruginosa* generally compromised *H. influenzae* growth (Figure 5). However, inhibition changed over time (AUC: linear mixed-effects model: F_2, 77_ = 9.55, p = 0.0002) and progressively declined towards a more neutral interaction (T1 vs. T2: estimate = –0.139 ± 0.077, *t*_77.4_ = –1.80, *p* = 0.1749; T1 vs. T3: estimate = – 0.347 ± 0.080, *t*_76.7_ = –4.34, *p* = 0.0001; T2 vs. T3: estimate = –0.208 ± 0.077, *t*_76.5_ = –2.72, *p* = 0.0218, Figure S6E). In turn, *H. influenzae* supernatant did not affect the overall growth (AUC) (Figure S6A) of *P. aeruginosa* but slightly prolonged the lag-phase (Figure 5, Figure S6B). This mild inhibitory effect changed over time (linear mixed-effects model: F_2, 297_ = 23.99, p < 0.0001) and became stronger (T1 vs. T2: estimate = 0.066 ± 0.001, *t*_72_ = 6.75, *p* < 0.0001; T1 vs. T3: estimate = 0.046 ± 0.001, *t*_72_ = 4.71, *p* < 0.0001; T2 vs. T3: estimate =-0.0201 ± 0.001, *t*_72_ =-2.05, *p* = 0.1081).
(ii) *S. aureus* versus *H. influenzae* – both species are early colonizers of the lungs and given their long-term persistence there is the potential for interaction patterns to change. We indeed observed that the effect of *S. aureus* supernatant on *H. influenzae* significantly changed over time (F_2, 70_ = 3.29, *p* = 0.0432), showing increasingly inhibitory effects (AUC, T1 vs. T3: estimate = 0.251 ± 0.098, *t*_72_ = 2.56, *p* = 0.0330; T1 vs. T2: estimate = 0.131 ± 0.098, *t*_72.1_ = 1.22, *p* = 0.3757; T2 vs. T3: estimate = 0.119 ± 0.098, *t*_72_ = 1.34, *p* = 0.4452, Figure 5, Figure S6E). Conversely, the effect of *H. influenzae* supernatant on *S. aureus* growth remained stable over time (linear mixed-effects model: F_2, 72_ = 2.37, *p* = 0.1004, Figure 5, Figure S6C).
(iii) *P. aeruginosa* versus *S. aureus* – Studies conducted with laboratory strains often reveal that *P. aeruginosa* is dominant over *S. aureus* ^44,45^ and that co-existence often relies on spatially structured environments where the two species are not well mixed ^46,47^. Accordingly, we expected that supernatants from *P. aeruginosa* isolates should inhibit the growth of *S. aureus* isolates. However, our data do not support this scenario. *P. aeruginosa* supernatant had positive effects on growth (AUC) (Figure S6C) and overall neutral effects on the lag phase of *S. aureus* (Figure 5, Figure S6D). Moreover, beneficial growth effects increased over time (AUC linear mixed-effects model: F_2, 72_ = 4.12, p = 0.0203; T1 vs. T2: estimate = 0.025 ± 0.149, t_72_ = 0.17, p = 0.9844; T1 vs. T3: estimate = 0.348 ± 0.155, t_72_ = 2.24, p = 0.0711; T2 vs. T3: estimate = 0.373 ± 0.140, t_72_ = 2.66, p = 0.0256) (Figure S6C). Similarly, lag-phases became shorter over time (F_2, 72_ = 13.26, p < 0.0001; T1 vs. T2: estimate =-0.023 ± 0.027, t_72_ =-0.87, p = 0.6637; T1 vs. T3: estimate = 0.101 ± 0.028, t_72_ = 3.64, p = 0.0015; T2 vs. T3: estimate = 0.124 ± 0.025, t_72_ = 4.96, p < 0.0001) (Figure S6D). *P. aeruginosa* overall growth was not affected by *S. aureus* supernatants and remained stable over time (AUC, linear mixed-effects model: F_2, 72_ = 1.36, *p* = 0.2632) (Figure S6A). In contrasts, lag phases were mildly negatively affected by *S. aureus* supernatants with lag phases becoming slightly longer over time (F_2, 72_ = 8.21, *p* = 0.0006; T1 vs. T2: estimate = 0.044 ± 0.011, *t*_72_ = 3.91, *p* = 0.0006; T1 vs. T3: estimate = 0.032 ± 0.011, *t*_72_ = 2.87, *p* = 0.0148; T2 vs. T3: estimate =-0.0117 ± 0.011, t =-1.05, *p* = 0.5485) (Figure S6B).

Altogether, our supernatant-based interaction assays indicate that, although *P. aeruginosa* becomes clinically dominant in CF lungs, it is not universally inhibitory to co-resident species under these conditions. *S. aureus* benefits overall from the presence of *P. aeruginosa* supernatant and the initial inhibition observed for *H. influenzae* was attenuated over time. *S. aureus* isolates, meanwhile, seem to become even stronger over time as supernatants from T3 have stronger negative effects on both *H. influenzae* and *P. aeruginosa*.

### Temporal shifts in species interactions are driven by a combination of donor and receiver effects

Our next goal was to determine whether the observed shifts in pairwise pathogen interaction patterns were driven by alterations in supernatant composition (donor effect) or by changes in the strains themselves (receiver effect). Donor effects manifest when supernatants have consistent growth effects on isolates regardless of the time window (T1, T2, T3) the isolates originate from. Receiver effects manifest when isolate growth is consistent regardless of the time window the supernatant originates from. Accordingly, we exposed all isolates to all supernatants across all three time windows and measured the corresponding growth effects (Figure S7). Using this approach and by focusing on overall growth (AUC), we uncovered four major patterns (Figure 6):

(i) *Temporal increase of H. influenzae growth in P. aeruginosa supernatant is driven by a donor effect*

**Figure 6.**
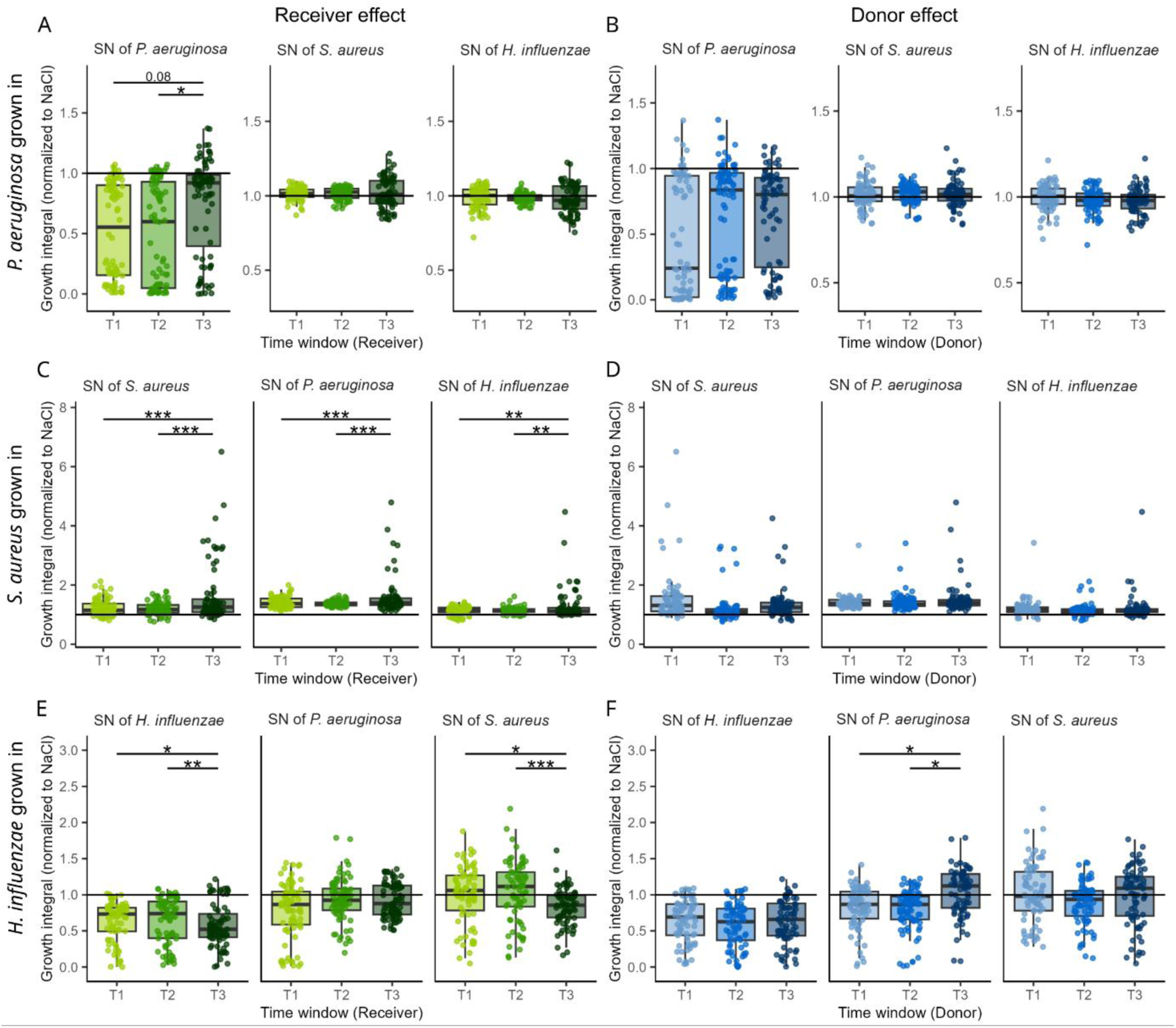
Across time window supernatant assays reveal one donor effect (reduced *P. aeruginosa* supernatant toxicity at T3) and several receiver effects (T3 isolates cope better with supernatants than isolates from earlier time windows). Green boxplots in panels (**A), (C)** and **(E)** show receiver effects for *P. aeruginosa*, *S. aureus* and *H. influenzae* isolates, respectively. Receiver effects were measured by exposing isolates from a specific time window T1, T2, or T3 to supernatants from respective donors (indicated on top of each panel) from all three time windows. Variation in growth across isolate time windows indicates that receivers differ and not the composition of the supernatants. Blue boxplots in panels **(B), (D)** and **(F)** show donor effects for *P. aeruginosa*, *S. aureus* and *H. influenzae* isolates, respectively. Donor effects were measured by exposing isolates from all time windows to supernatants sampled from a specific time window T1, T2, or T3. Variation in growth across supernatant time windows indicates that the composition of the supernatant (i.e., the donor) differs and not the receiving isolates. Receiver and donor effects were measured based on normalized growth (AUC relative to control condition) of isolates exposed to supernatants. We repeated the same analyses for the lag phase as an additional growth parameter (see Figure S8).

We found that the temporally decreasing inhibition of *H. influenzae* by *P. aeruginosa* supernatants was driven by a donor effect (linear mixed-effects model: F_2, 42_ = 5.38, p = 0.0083). Specifically, *P. aeruginosa* supernatants collected from T3 were no longer inhibitory against *H. influenzae* isolates, regardless of the time window the HI isolates originated from (Figure 6F, AUC comparisons: T1 vs. T3: estimate = 0.222 ± 0.083, *t*_42.2_ = –2.67, *p* = 0.0284; T2 vs. T3: estimate = 0.248 ± 0.083, *t*_41.8_ = 2.99, *p* = 0.0128; T1 vs. T2: estimate = 0.026 ± 0.083, *t*_42_ = 0.31, *p* = 0.9491). In contrast, there was no sign of a *H. influenzae* receiver effect (F_2, 42_ = 1.70, p = 0.1944). These results indicate that *P. aeruginosa* isolates from T3 reduced secretion of inhibitory factors compared to isolates from earlier time windows.

(ii) *Temporal decrease of H. influenzae growth in S. aureus supernatant is driven by a receiver effect*

We found that the temporally increasing inhibition of *H. influenzae* by *S. aureus* supernatants was driven by a receiver effect (linear mixed-effects model: F_2, 42_ = 8.80, *p* = 0.0006). Specifically, *H. influenzae* isolates from T3 grew significantly worse than isolates from T1 and T2 when exposed to *S. aureus* supernatants from all time windows (Figure 6E, AUC comparisons: T1 vs. T3: estimate = - 0.182 ± 0.060, *t*_42_ =-3.03, *p* = 0.0114; T3 vs. T2: estimate =-0.242 ± 0.060, *t*_42.1_ =-4.03, *p* = 0.0007; T1 vs. T2: estimate = 0.060 ± 0.060, *t*_41.9_ = –1.00, *p* = 0.5803). Conversely, there was no sign of a *S. aureus* donor effect (linear mixed-effects model: F_2, 42_ = 0.74, p = 0.4836). Identical receiver effects were observed for *H. influenzae* when exposed to supernatants from other isolates from its own species (AUC comparisons: T1 vs. T3: estimate =-0.082 ± 0.031, *t*_41.8_ =-2.63, *p* = 0.0310; T2 vs. T3: estimate = 0.104 ± 0.031, *t*_41.8_ = 3.35, *p* = 0.0048; T1 vs. T2: estimate =-0.022 ± 0.031, *t*_42.1_ =-0.71, *p* = 0.7579).

Altogether, these results indicate that *H. influenzae* isolates from T3 differ from earlier isolates by being more susceptible to compounds secreted by *S. aureus* and by isolates of its own species.

(iii) *Temporal increase of S. aureus growth in P. aeruginosa supernatant is driven by a receiver effect*

We found that the temporally increasing promotion of *S. aureus* by *P. aeruginosa* supernatants was driven by a receiver effect (linear mixed-effects model: F_2, 42_ = 12.95, p < 0.0001). Specifically, *S. aureus* isolates from T3 grew significantly better than isolates from T1 and T2 when exposed to *P. aeruginosa* supernatants from all time windows (Figure 6C, AUC comparisons: T1 vs. T3: estimate = 0.1945 ± 0.0481, *t*_42_ = 4.05, *p* = 0.0006, T2 vs. T3: 0.2257 ± 0.0481, *t*_42_ = 4.70, *p* = 0.0001, T1 vs. T2: estimate = 0.0312 ± 0.0481, *t*_42_ = 0.65, *p* = 0.7934). In contrast, there was no sign of a *P. aeruginosa* donor effect (Figure 6D, linear mixed-effects model: F_2, 42_ = 0.41, *p* = 0.6648). Identical receiver effects were observed for *S. aureus* when exposed to *H. influenzae* supernatants (Figure 6C, linear mixed-effects model: F_2, 42_ = 8.60, p = 0.0007) and supernatants from other isolates from its own species (Figure 6C, linear mixed-effects model: F_2, 42_ = 11.77, p < 0.0001). Taken together, these results suggest that *S. aureus* isolates from T3 differ from earlier isolates by being better at growing in spent supernatant medium.

(iv) *Temporal decrease of inhibition among P. aeruginosa isolates is driven by a receiver effect*

We found that temporal changes in the inhibition that *P. aeruginosa* isolates had on each other were driven by a receiver effect (Figure 6A, linear mixed-effects model: F_2, 42_ = 3.86, *p* = 0.0288). Specifically, *P. aeruginosa* strains isolated from T3 exhibited significantly higher growth regardless of the time window the supernatant originated from (AUC comparisons: T1 vs. T3: estimate = 0.215 ± 0.098, *t*_42_ = 2.20, *p* = 0.0830; T2 vs. T3: estimate = 0.251 ± 0.098, *t*_42_ = 2.57, *p* = 0.0359; T1 vs. T2: estimate = 0.037 ± 0.098, *t*_42_ = 0.37, *p* = 0.9264).

### Reduction of protease activity among *P. aeruginosa* isolates is associated with improved growth of *S. aureus* and *H. influenzae* isolates

The above analyses revealed a single donor and several receiver effects. Here, we focus on the donor effect of *P. aeruginosa* supernatant on *S. aureus* and *H. influenzae* growth with the aim of establishing a functional link with the observed phenotypic changes in *P. aeruginosa*. For this we built a generalized additive model (GAM) by either fitting mean normalized AUC or lag phase as response variable and the phenotypes either as factor (hemolysis status) or co-variates (smooth terms, biofilm formation, siderophore and protease production).

For *S. aureus*, the lag phase model had the highest predictive value and explained 27.7% of the deviance (adjusted R² = 0.26). Higher protease production in *P. aeruginosa* was significantly and non-linearly associated with prolonged lag phases in *S. aureus* (Figure 7A; edf = 3.51, F = 9.74, p < 0.0001) and hemolytic *P. aeruginosa* isolates inhibited *S. aureus* more strongly than non-hemolytic ones (β = −0.060 ± 0.0289, t = −2.08, p = 0.0387). The other *P. aeruginosa* phenotypes were not significantly associated with the *S. aureus* lag-phase (Figure S9, biofilm formation: edf = 1.00, F = 1.16, p = 0.2834; siderophore production: edf = 1.00, F = 0.84, p = 0.3595).

**Figure 7.**
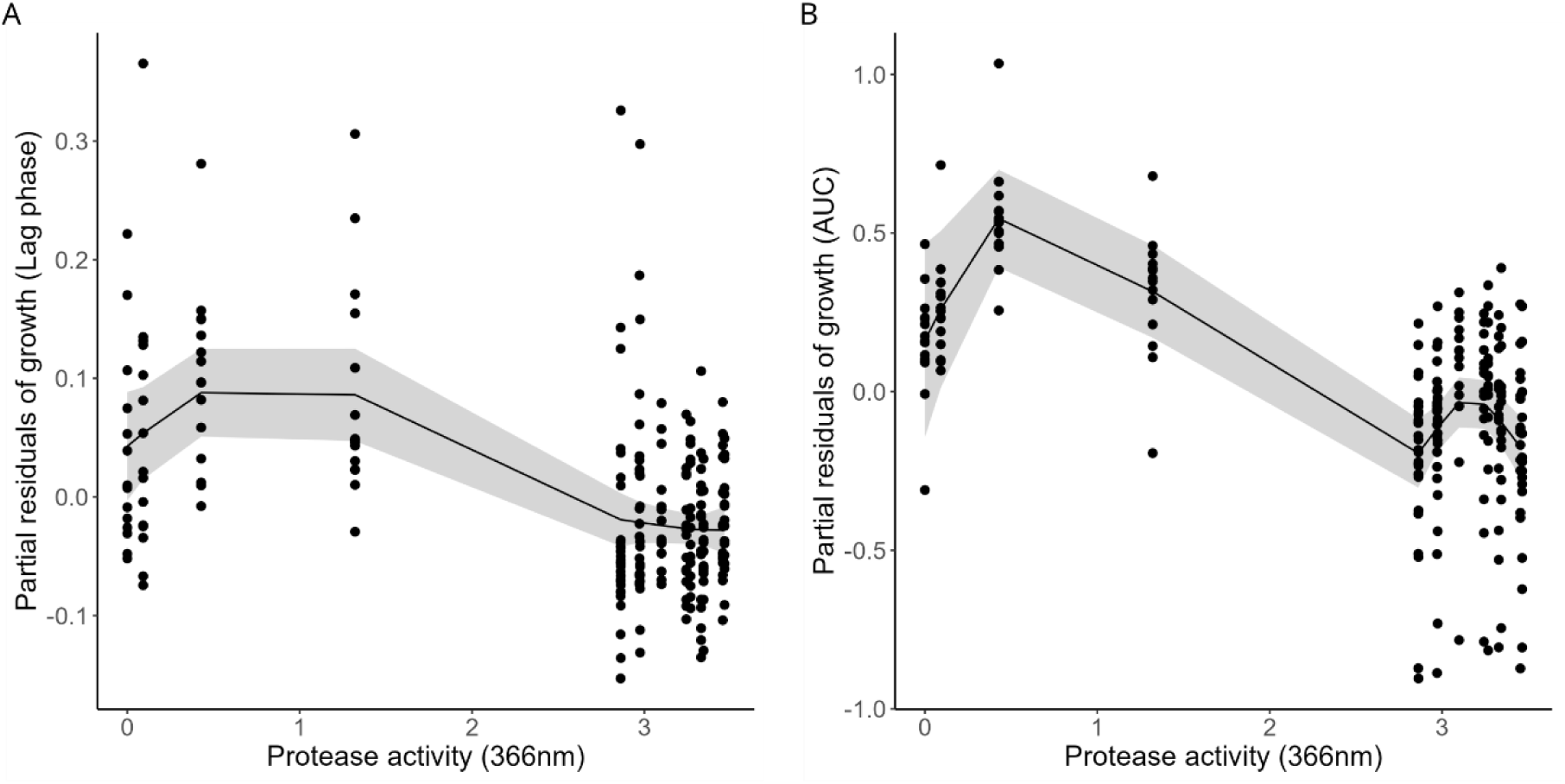
Effect of protease production in *P. aeruginosa* on bacterial growth of *S. aureus* and *H. influenzae* estimated from generalized additive models (GAMs). (A) Relationship for *S. aureus* based on normalized lag phase data (log-transformed). **(B)** Relationship for *H. influenzae* based on normalized AUC (area under the curve) data. Both associations show the smoothed effect of protease activity with 95% confidence intervals (shaded ribbons), with points representing partial growth residuals. GAM models included hemolysis as categorical predictor and smooth terms for log-transformed biofilm formation, siderophore and protease production.

For *H. influenzae*, the AUC model had the highest predictive value and explained 44.9% of the deviance (adjusted R² = 0.43). We found that higher protease production in *P. aeruginosa* was associated with significantly stronger *H. influenzae* inhibition (Figure 7B; edf = 5.13, F = 13.02, p < 0.0001). The other *P. aeruginosa* phenotypes were not significantly associated with *H. influenzae* growth (siderophore production: edf = 1.69, F = 0.66, p = 0.5536; biofilm formation: edf = 1.00, F = 2.51, p = 0.1150; hemolysis (β = −0.018, t = −0.143, p = 0.8860).

Altogether, these results suggest that a reduction of protease activity among T3 isolates of *P. aeruginosa* is the main variable responsible for improved growth of both *S. aureus* (lag-phase) and *H.influenzae* (AUC). Protease production is mainly controlled by the Las quorum-sensing system, and mutations in this system are often associated with reduced protease activity ^48^. Consistent with these findings, we detected missense point mutations and deletions, both of which can lead to loss of function ^48^, in all key genes of the Las quorum-sensing system (*lasI* signal synthase, *lasR* receptor, and *lasB* protease) ^49^ relative to the reference *P. aeruginosa* PAO1 in isolates with reduced protease activity (Table S3; Figure S10; ID114: *lasI*, ID115: *lasB*, ID116: *lasR*, ID117: *lasR*).

## Discussion

Cystic fibrosis (CF) is a life-limiting hereditary disease in which thick airway mucus creates nutrient-rich niches that promote lung colonization by different pathogens ^50,51^. Infections in CF lungs are typically polymicrobial following an ecological succession, whereby early colonizers such as *H. influenzae* and *S. aureus* are followed by late colonizers such as *P. aeruginosa* ^2,52,53^. Upon colonization of the complex lung environment, pathogen can adapt to various components, including host factors, other pathogens but also medical treatment. Moreover, since the lung is an open system new strains can invade over time potentially leading to ecological turnovers ^54^. While such ecological and evolutionary dynamics have been intensively studied in lungs of adult people with CF for individual pathogen species, little is known about pathogen dynamics and species interactions during early childhood. To approach this knowledge gap, we analyzed longitudinally sampled clinical isolates from 23 children with CF (0–8 years of age) comprising the three most abundant species *P. aeruginosa*, *S. aureus*, and *H. influenzae* ^27^. By combining whole-genome sequencing, phenotypic assays, and supernatant interaction experiments, we found considerable temporal variation in genome composition, virulence-associated phenotypes and interspecies interaction patterns. Our data suggest that both ecological turnover of pathogen variants and the persistence and evolution of pathogens within children with CF occur. When focusing on interspecies interactions, we found that the most prominent change involved *P. aeruginosa,* whereby isolates from older children (> 5 years of age) frequently carried mutations in quorum sensing (QS) genes. These isolates exhibited reduced protease production, which correlated with attenuated inhibition of *S. aureus* and *H. influenzae* in supernatant assays. This shows how a single genetic change in one species can affect the interspecies interaction network, shifting it toward less antagonistic relationships.

The main goal of our study was to examine ecological and evolutionary patterns within and between members of the pathobiome colonizing the lungs of children with CF. However, several of our analyses need to be interpreted with care as the temporally heterogenous and non-systematic sampling of isolates reflects a limitation and prevents the precise tracking of within-individual ecological and evolutionary dynamics. Instead, we had to pool all isolates across children and assume that eco-evolutionary conditions are similar across individuals. Despite this limitation, several results indicate that a combination of ecological and evolutionary factors jointly influence pathobiome dynamics. First, among the children from which multiple consecutive samples were available we found significant phylogenetic clustering among isolates in several individuals (for *P. aeruginosa*: P6, P15; for *S. aureus*: P5, P6, P9, P14). These isolates shared the same serotype and *spa* type, respectively, suggesting that the same isolates persist over time, a prerequisite for evolution to enact. Second, we observed the accumulation of *P. aeruginosa* isolates with mutations in quorum sensing (QS) regulators among samples from the last time window (T3). These mutations were all associated with reduced protease production. The accumulation of QS-mutations has repeatedly been shown to be a hallmark of evolutionary adaptation to the CF lung environment in adult people with CF ^55–57^. The observation that such QS mutations surface in children with CF is an indicator that adaptive evolution is already operating in young children. Third, we found clear evidence for ecological strain turnover in several individuals (for *S. aureus*: P3, P22; for *H. influenzae*: P8, P10, P12), whereby earlier isolates were replaced by new isolates belonging to different phylogenetic clusters (and *spa* types for *S. aureus*). Altogether, these results suggest a mixed model for the pediatric lung environment characterized by both pathogen isolate turnover and longer-term persistence of specific isolates.

The lung is a complex environment, and it is notoriously hard to disentangle correlative patterns from causes and consequences. In the context of interactions between different members of the pathobiome, we would like to highlight four scenarios that may result in a change in interactions pattern as a by-product and not as a result of adaptations of pathogens towards each other. First, lung physiology and host immune status (e.g., inflammation) change during childhood. These environmental changes may determine the bacterial genotypes and phenotypes that can invade the lung eco-system and spur successive strain turnover for each pathogen species ^58,59^. Strain turnover can lead to changes in species interaction patterns without adaptive evolution being involved. Second, each pathogen species may independently adapt to the lung environment, with these adaptations indirectly feeding back on pathogen interaction patterns. For example, the lung environment could select for strains with increased biofilm formation, which in turn can reduce competitive interactions between species ^15^. Third, *P. aeruginosa* is known to diversify within the lung environment ^14,60^. Diversification can favor adaptation to different niches but also foster competition between lineages. For instance, lineages that are deficient to produce proteases and siderophores can enjoy relative fitness advantages by exploiting the molecules secreted by producer lineages ^21,61^. Such within-species competitive dynamics can affect between-species competitive interactions. Fourth, children with cystic fibrosis commonly receive antibiotics targeting *S. aureus* and *H. influenzae*, and specific therapy against *P. aeruginosa* upon detection of this pathogen. Evolutionary responses to treatment may alter pathogen interaction patterns. For instance, selection for resistance mechanisms in *P. aeruginosa* could result in cross protection of non-targeted bystander species such as *S. aureus* and *H. influenzae*^23^. In reality, changes in pathogen interactions, like those observed in our study, are probably the result of both by-products of adaptations to the above-mentioned factors and direct adaptation to co-infecting species.

From a community perspective, it is important to ask whether changes in interaction patterns between two pathogens leads to changes in community stability. There are at least three possible scenarios. First, one species could gain dominance over another one and displace it from the community. For instance, our findings suggest that *H. influenzae* gets increasingly suppressed by *S. aureus* over time and might go extinct in the long run. Second, competition between species may attenuate over time, leading to more stable co-existence. Reduced inhibition of *H. influenzae* by *P. aeruginosa* over time might be an example of this scenario. Third, species could engage in antagonistic co-evolution, characterized by repeated adaptation and counteradaptations by each of the two species against the opponent. Our limited data set does not allow to assess the relative importance of the three scenarios. But we anticipate that strong antagonistic co-evolution may be the least likely scenario, as not all species are present in the lung all the time and they may often occupy different niches and thus not regularly interact. Furthermore, it would be important to test whether changes in interactions between different pairs of pathogens are additive or non-additive. For example, it would be interesting to test whether the increasingly negative interactions *H. influenzae* experiences against *S. aureus* are offset by the more positive interactions it has with *P. aeruginosa*.

In summary, we studied genetic and phenotypic variation across three key species of the pathobiome of children with CF and related the observed variation to ecological changes in interaction networks between pathogens. We show that genotypes and phenotypes typically linked to chronic CF infection can already be found at a young age. Notably quorum-sensing and siderophore deficient isolates are common among *P. aeruginosa* isolates from adult people with CF with such isolates often showing attenuated virulence and altered community interactions ^55^. Similarly, *S. aureus* small-colony variants are also frequently isolated from adult people with CF and have been associated with persistence, intracellular survival, and poorer clinical outcomes ^62–64^. Conversely, mucoid phenotypes that are characteristic for *P. aeruginosa* isolates from adult people with CF are absent among our pediatric isolates. Moreover, our data suggest that the pathobiome is characterized by a certain degree of strain turnover, which is less commonly observed in chronic infections of adults ^65^. A key question for future research is to assess changes in pathogen interaction patterns at the level of the individual pathogen and how such changes influence host health and disease progression. Overall, the understanding of early patterns of lung colonization, pathogen trait variation, and changes in interactions among pathobiome members may prove useful for deciding on treatments that maintain benign communities and delay the establishment of chronic, hard-to-treat infections.

## Material and Methods

### Bacterial strains and culture conditions

We used clinical *Pseudomonas aeruginosa* (19), *Staphylococcus aureus* (44) and *Haemophilus influenzae* (21) strains that were isolated from oropharyngeal swap samples from children with CF, who are part of the SCILD cohort (https://www.scild.ch) (Table S1) ^27^. Samples were taken by Prof. Dr.

Philipp Latzin and colleagues, and the strains were isolated by Dr. Markus Hilty. Furthermore, we used laboratory strains, including *P. aeruginosa* PAO1 (ATCC 15692), *S. aureus* strains USA300 JE2 (ATCC-BAA-1717), Cowan 1 (ATCC 12598), and 6850 (ATCC 53657), and *H. influenzae* strains ATCC 49766 (NTHI), ATCC 49247 (NTHI) and ATCC 10211 (Serovar B) (Table S4).

For all assays, we used BHI medium supplemented with the following stock solutions: 10ml β-nicotinamide adenine dinucleotide (β-NAD; 1.51 mM), 10 ml hemin [1.53 mM], 4.86 ml of MgCl2 [1 M] and 1.6 ml CaCl2 [1 M]. Hereafter called ssBHI. We stored all strains in ssBHI containing 40% glycerol at-80 degrees Celsius. We grew overnight cultures in 4 mL ssBHI within 13 mL tubes at 37 degrees Celsius and 170 rpm. The next day, we washed overnight cultures with 0.8% sodium chloride (NaCl) solution and adjusted them to an optical density at 600 nm (OD_600_) of 0.0004.

### Genome sequencing

We individually harvested cells of all our isolates and resuspended them in a tube with DNA/RNA Shield (Zymo Research, USA). 5 - 40 μl of the cell suspension are lysed with 120 μL of TE buffer containing lysozyme (MPBio, USA) metapolyzyme (Sigma Aldrich, USA) and RNase A (ITW Reagents, Spain), incubated for 25 minutes at 37 degrees Celsius. Proteinase K (VWR Chemicals, Ohio, USA) (final concentration 0.1mg/mL) and SDS (Sigma-Aldrich, Missouri, USA) (final concentration 0.5% v/v) are added and incubated for 5 minutes at 65 degrees Celsius. Genomic DNA is purified using an equal volume of SPRI beads and resuspended in EB buffer (10mM Tris-HCl, pH 8.0).

Extracted DNA is then quantified with the Quant-iT dsDNA HS (ThermoFisher Scientific) assay in an Eppendorf AF2200 plate reader (Eppendorf UK Ltd, United Kingdom) and diluted as appropriate.

Genomic DNA libraries are prepared using the Nextera XT Library Prep Kit (Illumina, San Diego, USA) following the manufacturer’s protocol with the following modifications: input DNA is increased 2-fold, and PCR elongation time is increased to 45 seconds. DNA quantification and library preparation are carried out on a Hamilton Microlab STAR automated liquid handling system (Hamilton Bonaduz AG, Switzerland). Libraries are sequenced on an lllumina NovaSeq 6000 (Illumina, San Diego, USA) using a 250 bp paired end protocol. Reads are adapter trimmed using Trimmomatic version 0.30 ^66^ with a sliding window quality cutoff of Q15. De novo assembly is performed on isolates using SPAdes version 3.7 ^67^, and contigs are annotated using Prokka 1.11 ^68^.

### Genome analysis

A pangenome was constructed with Roary (v3.13.0) ^69^, for (i) a core-gene alignment for phylogenetic inference and (ii) a binary gene presence–absence matrix for gene-content analyses. A species phylogeny was inferred with OrthoFinder (v2.5.5) ^70,71^, which reconstructs orthogroups, infers individual gene trees with FastTree (v2.1.10) ^72^, and estimates a consensus species tree using the STAG^73^ algorithm with rooting performed by STRIDE ^74^. Gene families were assigned to frequency bins defined as core (present in 100% of strains), soft core (present in ≥90% and <100%), shell (present in ≥15% and <90%), and cloud (present in <15%); bins were treated as mutually exclusive. Protein sequences were functionally annotated with eggNOG-mapper (v2.1.12) ^75^using the eggNOG 5.0.2 database to assign COG functional categories ^30^. Where a gene mapped to multiple categories, all assigned categories were retained for counts and enrichment summaries. Putative virulence loci were identified by searching predicted proteins against the VFDB (core dataset, release May 2024) ^76^ using using DIAMOND/BLASTP (v2.1.10) with standard stringency (e-value ≤ 1 × 10⁻⁵, percent identity ≥ 90%, alignment coverage ≥ 80%). Sequences for each locus were collated and, when required, aligned with MAFFT (v7.525; L-INS-i mode) for comparative analyses. Genome-wide gene-content dissimilarity was quantified as Jaccard distance computed from the Roary presence–absence matrix using R (v4.3) and the vegan package, and distances were summarized within and between children with CF time windows.

### Hemolysis assay

To evaluate the hemolytic capabilities of our isolates, we cultured all isolates overnight in ssBHI and then plated them on COS plates (Columbia agar supplemented with 5% sheep blood, Biomérieux, Petit-Lancy, Switzerland). We incubated the COS plates anaerobically with 5% CO_2_ at 37 degrees Celsius overnight. The following day, we qualitatively assessed the hemolysis patterns of each isolate based on their ability to lyse the red blood cells. Hemolytic activity was recorded using a qualitative yes or no classification.

### Small colony variant detection

*Staphylococcus aureus* isolates were cultured on mannitol salt agar at 37 degrees Celsius in 5% CO₂ for 48 hours. Plates were examined for the presence of small colony variants (SCVs), which were defined as colonies with a diameter <50% of that of typical *S. aureus* colonies on the same plate.

### Mucoidy detection

To assess the mucoidy of *Pseudomonas aeruginosa* isolates, cultures were directly plated from cryo stocks onto solidified supplemented brain heart infusion (ssBHI) agar plates. Plates were incubated at 37 degrees Celsius for 24 hours. Following incubation, colony morphology was evaluated using a sterile inoculation loop. Isolates were classified as mucoid if they exhibited slimy, viscous growth when gently touched with the loop. Mucoidy was assessed qualitatively yes or no classification. The laboratory strain PAO1 was included as a non-mucoid reference control.

### Protease production quantification

To assess protease production in the clinical isolates from all three species, we conducted the azocasein assay to determine their protease activity, adapting the protocol from Chessa *et al.* (2000)^77^. We acknowledge however, that this assay may not capture all proteases produced by the isolates, such as some secreted by *S. aureus*. We grew the isolates overnight in ssBHI medium and re-suspended them to a final OD_600_ of 0.01 before incubating them in 96-deep well plates with 1.5 ml ssBHI for 22 hours at 37 degrees Celsius and 170rpm. The next day, we transferred 200 μl to 96 well plates to measure the growth in a plate reader. Afterwards, we centrifuged the plates for 30 minutes at 3700 rpm to pellet cells and transferred 200 μl of the cell-free supernatants to fresh 96-well plates before storing them in the freezer at-20 degrees Celsius. Next, we treated 40 μl of thawed aliquots with 120 μl phosphate buffer (50 mM [∼7.5 pH]) and 40 μl azocasein (30mg/ml) and incubated the mixture for 20 hours at 37 degrees Celsius under static condition. We added 200 μl trichloroacetic acid (20%) to stop the reaction and centrifuged the treated supernatants at 3700 rpm for 15 minutes. Finally, we collected and transferred the fresh supernatants into new 96-well plates and quantified the protease activity as optical density in a plate reader at 366nm. Blank ssBHI medium treated the same as the supernatants was used as a blank control. We repeated the experiment three times.

### Siderophore production assay

To evaluate whether the clinical isolates can produce pyoverdine, we grew all isolates under iron-limited conditions and assessed their siderophore activity using the chrome azurol S (CAS) assay to measure pyoverdine production levels. To acquire this, we adjusted the protocol from Sathe *et al.* (2019) ^78,79^. We prepared overnight cultures in 24-well plates with 1.5 ml ssBHI growth media without adding hemin for *P. aeruginosa* and *S. aureus* and incubated them for 20 hours at 37 degrees Celsius and 170rpm. Then, we adjusted the OD_600_ to 0.01 and incubated the strains as triplicates in 24-well plates with 1.5 ml ssBHI supplemented 2,2’-bipyridine to a final concentration of 600 μM (*P. aeruginosa*) and 75 μM (*S. aureus* and *H. influenzae*) respectively to ensure siderophore production for 24 hours at 37 degrees Celsius and 170 rpm. After 24 hours, the final OD_600_ was measured to assess the growth. We then poured the supernatants of the triplicates together, centrifuged for 10 minutes at 7000 rpm before filter sterilizing (0.2-μm pore size, Whatman) to get the prepared supernatant. To assess the siderophore activity, we mixed 100 μl of the sterile supernatant with 100 μl CAS solution in a 96-well plate and incubated for 20 minutes in the dark at room temperature. After the inoculation time, we measured the optical density at 630nm (OD_630_) in the microplate reader and calculated the “percentage of siderophore units” (psu). We repeated the experiment three times.

### Biofilm formation assay

We quantified surface-attached biofilms by a crystal violet assay. Single colonies were grown overnight from glycerol stocks in ssBHI (4 ml, 13 ml tubes, 18 hours, 37 degrees Celsius, 170 rpm), centrifuged (7000 rpm, 5 minutes, RT), washed in fresh ssBHI, and adjusted to OD_600_ = 0.01. Aliquots (200 µl) were inoculated into flat-bottom 96-well plates and incubated statically at 37 degrees Celsius for 24 hours (*P. aeruginosa*, *S. aureus*) or 48 hours (*H. influenzae*). Cultures were transferred to a fresh plate to read OD_600_ (growth control). Biofilms were washed with ddH2O, stained with 0.1% crystal violet (175 µl, 30 minutes, RT), washed ×3, air-dried (45 minutes), and destained with DMSO (200 µl, ∼20 minutes, RT). OD_570_ was measured and biofilm formation expressed as Biofilm Index = OD_570_/OD_600_ ^80^. ssBHI stained with crystal violet/DMSO served as the blank control. Experiments were performed in triplicate.

### Supernatant assay

To evaluate whether strains inhibit or promote the growth of other isolates, we performed a supernatant assay. Initially, we inoculated isolates from frozen glycerol stocks into 4 ml of ssBHI in 13 ml culture tubes, incubating them at 37 degrees Celsius and 170 rpm for 18 hours. The following day, we centrifuged the cultures for 5 minutes at 3700 rpm at room temperature, washed the cells with 0.9% NaCl, and adjusted the cell density to 6.65 x 10^5^ cells/ml. We then incubated these adjusted cells in 4 ml of ssBHI in 13 ml culture tubes, with six replicates per isolate, for 20 hours under the same conditions.

Post-incubation, we centrifuged the cells again for 5 minutes at 3700 rpm, combined the supernatants from the replicates, and filter-sterilized them using a 0.2-μm Whatman filter.

Concurrently, we washed one set of sextuplicate cultures with 6 ml of fresh 0.9% NaCl and adjusted the cell density to 6.65 x 10^5^ cells/ml. For the assay, we inoculated these washed and adjusted cells into a 200 μl mixture containing 125 μl of ssBHI and 75 μl of the prepared supernatant in 96-well plates, incubating them at 37 degrees Celsius and 170 rpm for 24 hours. We monitored the optical density at 600 nm (OD_600_) every 2 hours for the first 12 hours, then every 3 hours for the remaining time. As controls, we used a mixture of 125 μl ssBHI and 75 μl 0.9% NaCl inoculated with cells as a positive minimal control, 200 μl ssBHI as a positive maximal control, and blank ssBHI medium as a negative control.

### Determination of antibiotic dose-response curves

To test the clinical isolates for antibiotic resistance or susceptibility, we used the broth microdilution method in ssBHI with the following antibiotics: ampicillin, ciprofloxacin, colistin, meropenem, piperacillin–tazobactam (2:1), and tobramycin (Table S2). Antibiotic serial dilutions were prepared in ssBHI in sterile 96 deep-well plates, foil-sealed, and stored overnight at 4 degrees Celsius. Immediately before the assay, 100 µL from each well was transferred to 96-well assay plates using the MagPie multi-dispenser robot. In parallel, 1 mL of each overnight culture was transferred to 2 mL Eppendorf tubes, washed once with 1 mL 0.9% NaCl (6000 rpm, 5 minutes), and adjusted to ∼10⁶ cells/mL via two 1:10 dilutions. Then, 20 µL of the OD-adjusted bacterial suspension was dispensed into wells pre-filled with 180 µL antibiotic solution (final volume 200 µL). Columns 1–11 received inoculum, and column 12 contained 200 µL ssBHI only as a sterility control and for blank correction. An initial inoculation OD₆₀₀ reading was taken, plates were incubated for 24 hours at 37 degrees Celsius with shaking at 170 rpm, and endpoint 24 hours OD_600_ measurements were recorded.

### Statistical analysis

To assess changes in growth over time, we first compared normalized AUC values from combinations of strains and supernatants isolated at the same time window (T1, T2, or T3), using a linear mixed-effects model with time window as a fixed effect and the strain ID and supernatant ID as a random effect (Figure 5; Figure S6). To determine a receiver (strain-driven) from donor (supernatant-driven) effects, we fit two linear mixed-effects models. For the receiver effect (Figure 6A, C,E; Figure S7; Figure S8A,C,E), we analyzed growth and lag phase of strains from each time point across all supernatants, treating the receiver strain’s time window as a fixed effect and including a random intercept for supernatant ID nested within that time window. For the donor effect (Figure 6B, D, F; Figure S7; Figure S8B,D,F), we analyzed growth and lag phase in supernatants from each time point across all strains, treating the donor (supernatant) time window as a fixed effect and including a random intercept for receiver strain ID nested within the donor time window. All models were fit using the lmer function from the lme4 package in R.

To explore nonlinear effects of phenotypic traits, we fitted generalized additive models (GAMs) with the mgcv package in R. We analyzed two responses. First, normalized AUC for *H. influenzae* and second, the log-transformed normalized lag phase for *S. aureus*. Parametric term included hemolysis (binary), and nonlinear effects were modeled with smooth terms for log-transformed biofilm, siderophore production, and protease activity (Figure 7; Figure S9). Smoothing parameters were selected by generalized cross-validation (GCV) using the default option.

## Data availability

All other data supporting the findings of this study are available from the corresponding author upon request and will be made publicly available upon publication.

## Acknowledgments

This project was supported by funding from the Swiss National Science Foundation (grants no 212266 to R.Kü.), the University of Zurich Research Priority Program (URPP) Evolution in Action (to L.S.).

P.D.C.S. was funded by a PhD studentship from the Biotechnology and Biological Sciences Research Council Midlands Integrative Biosciences Training Partnership (grant reference BB/M01116X/1)

## Authors Contributions

Conceptualization was performed by L.S., F.H., M.H. and R.Kü. Sampling and isolation of bacterial strains were performed by M.H. and P.L. Data curation was performed by L.S. and R.Kü. Formal analysis was performed by L.S., R.Ku. and S.P. Investigation was performed by L.S., R.Ku., S.P., and P.D.C.S. Visualization was performed by L.S. Writing—original draft was performed by L.S. and R.Kü. Writing—review and editing was performed by all authors. All authors read and approved the final manuscript.

