## Supplemental Figures and Tables for "Tracking eco-evolutionary dynamics among lung pathobiome members from children with cystic fibrosis"

This document contains the following supplementary material:

10 Supplementary Figures

4 Supplementary Tables

**Supplemental Figure S1**

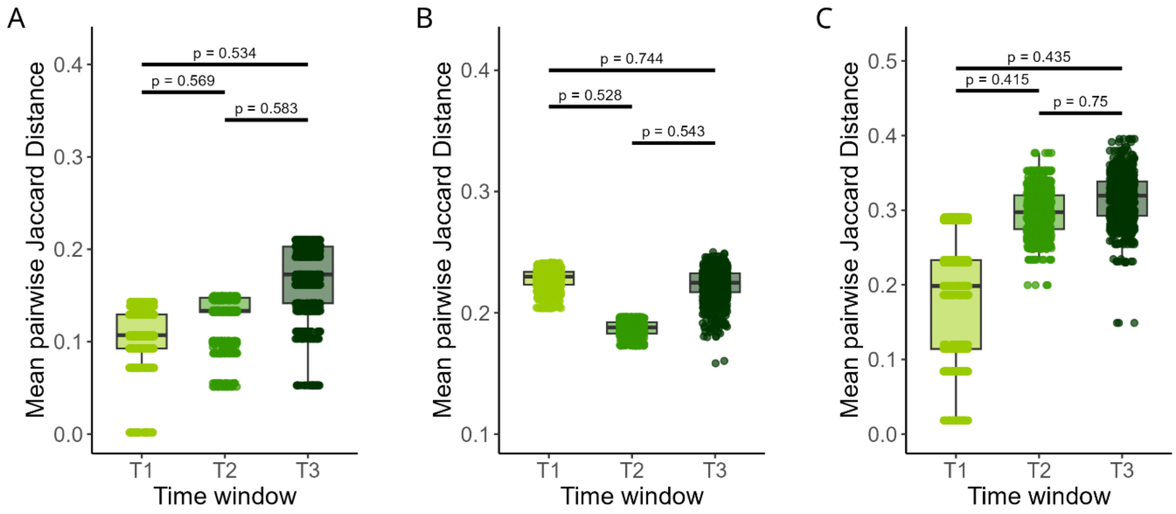


**Figure S1 | Within-time window Jaccard distance across three species.** Mean pairwise Jaccard distances were calculated within each time window (T1, T2, T3) for each species independently, based on Roary gene presence/absence matrices. Boxplots show the distribution of bootstrapped mean Jaccard distances across 1,000 iterations. Higher values indicate greater genetic dissimilarity among strains sampled within the same time window. Statistical comparisons between timepoints were performed using two-tailed permutation tests, with p-values annotated above the relevant groups. **(A)** Shows results for *Pseudomonas aeruginosa*, **(B)** for *Staphylococcus aureus* and **(C)** for *Haemophilus influenzae*.

### Supplemental Figure S2

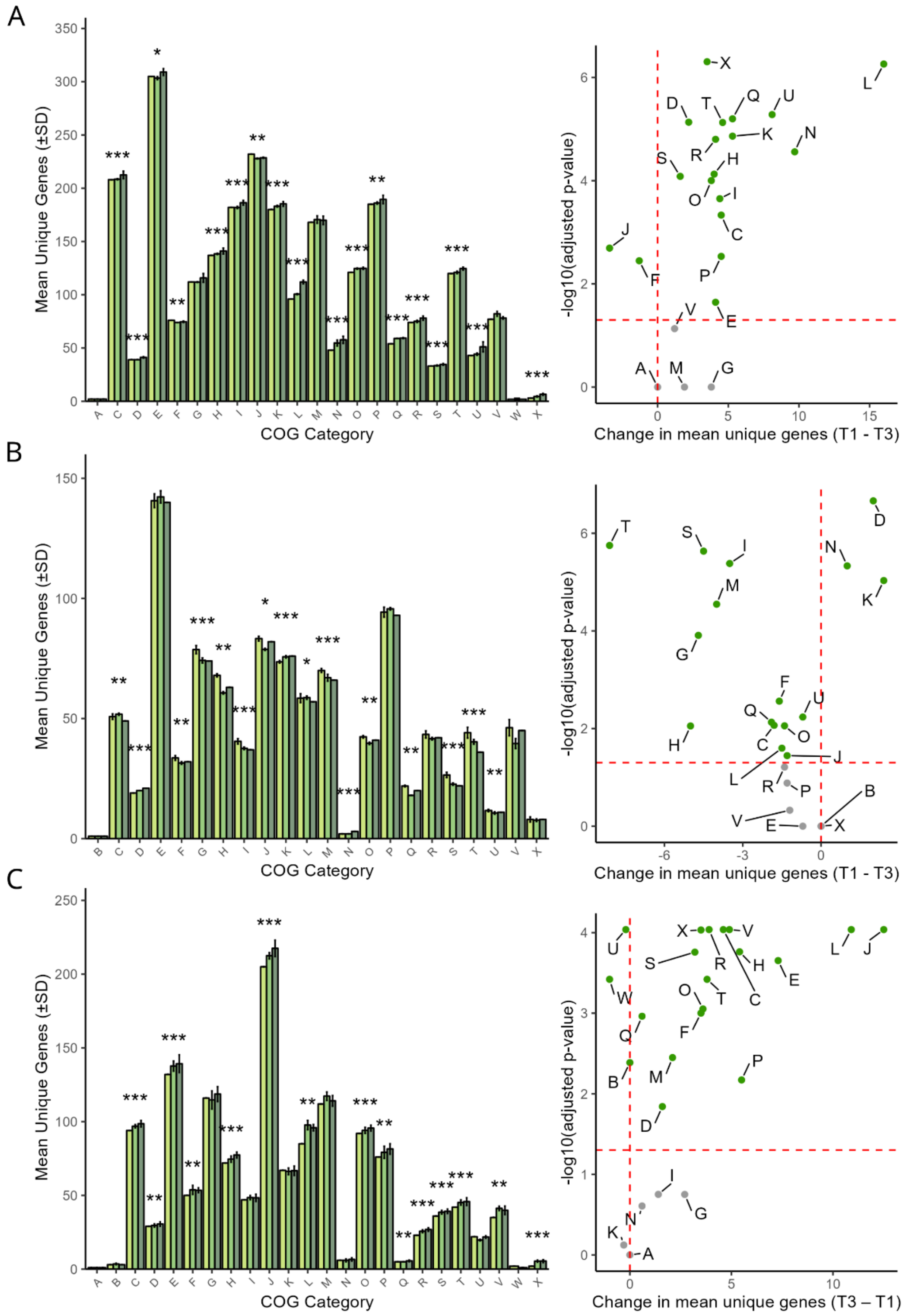

**Figure S2 | Temporal changes of COG categories across the three time windows.** Left panels show barplots showing the mean number of unique genes per COG category across time windows (T1, T2, T3), based on rarefied subsets of strains per group (n = smallest group size). Error bars represent standard deviations from 10 rarefaction iterations. Asterisks above bars indicate COG categories with significant differences between T1 and T3 (Dunn's post-hoc test, BH-adjusted $p < 0.05$). Left panels show volcano plots illustrating the difference in mean unique gene counts (T1 – T3) per COG category (x-axis) versus $-\log_{10}$ of the adjusted p-value (y-axis). Red dashed lines indicate the significance threshold ($p = 0.05$) and no-change baseline. COG categories with statistically significant differences are highlighted in green. **(A)** Shows results for *Pseudomonas aeruginosa*, **(B)** for *Staphylococcus aureus* and **(C)** for *Haemophilus influenzae*.

### Supplemental Figure S3

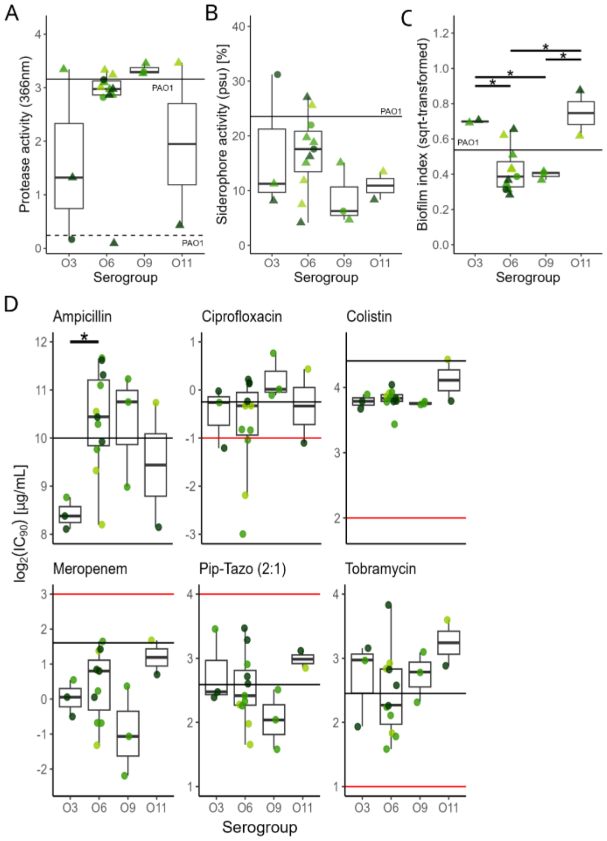


**Figure S3 | Phenotypes and $IC_{90}$ values of 19 *P. aeruginosa* isolates grouped by serogroup. (A)** Protease activity of clinical isolates measured with the azocasein assay. **(B)** Siderophore production of the clinical isolates measured with the chrome azurol S (CAS) assay in ssBHI medium supplemented with 600 µM 2,2'-bipyridine. **(C)** Biofilm production under static growth conditions measured with the crystal violate assay. **(D)** $IC_{90}$ values were determined in ssBHI medium for six antibiotics (ampicillin, ciprofloxacin, colistin, meropenem, piperacillin–tazobactam, and tobramycin) for *P. aeruginosa.* *P < 0.05.

**Supplemental Figure S4**

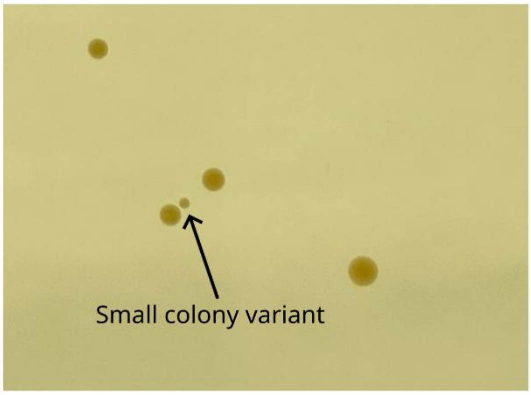


**Figure S4 | Small colony variant of *S. aureus* clinical isolate grown on mannitol salt agar.** The image shows a single small colony variant (SCV; presence of colonies with a diameter <50% that of the typical *S. aureus* diameter, indicated by the arrow) of *S. aureus* isolate ID228, alongside four colonies exhibiting normal morphology.

### Supplemental Figure S5

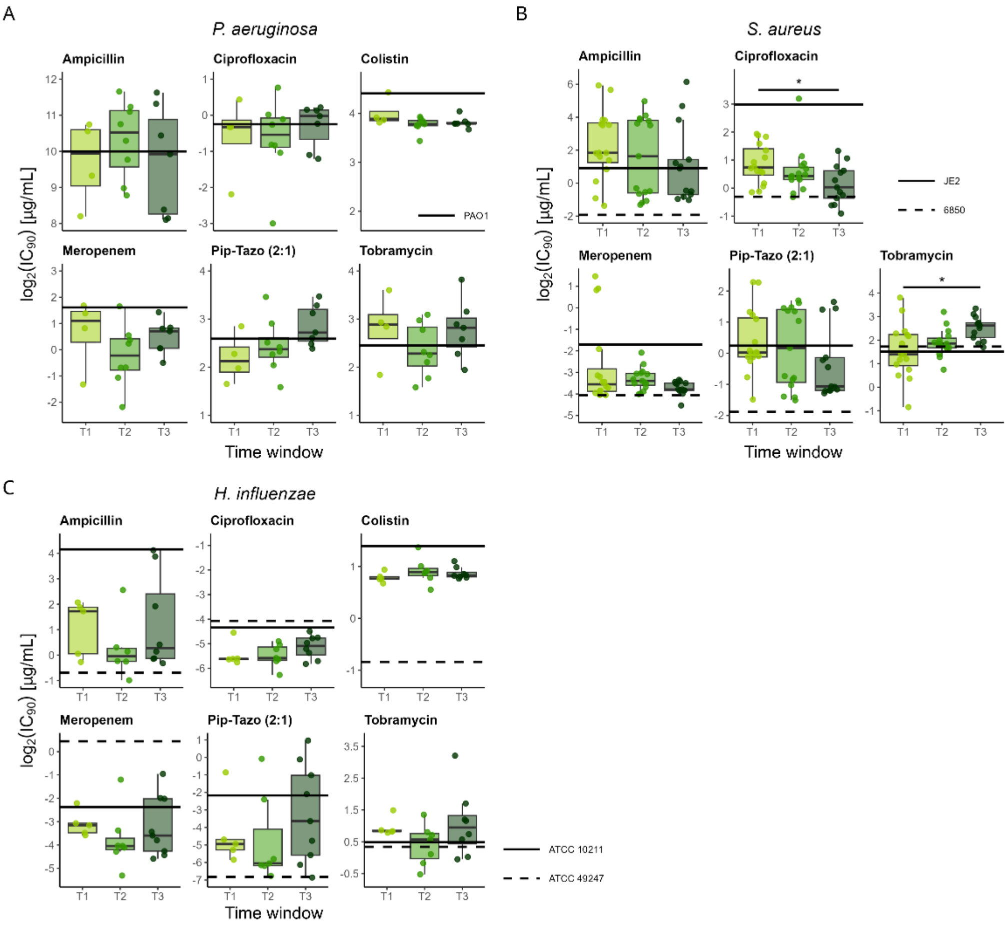


**Figure S5 | $Log_2$ transformed $IC_{90}$ values of the clinical isolates.** $IC_{90}$ values were determined in ssBHI medium for six antibiotics (ampicillin, ciprofloxacin, colistin, meropenem, piperacillin–tazobactam, and tobramycin) in **(A)** *P. aeruginosa* and **(C)** *H. influenzae*. For **(B)** *S. aureus*, only five antibiotics were tested (colistin was not included). Black lines (solid and dashed) represent the reference laboratory strains. Asterisks denote significance levels: *P < 0.05.

### Supplemental Figure S6

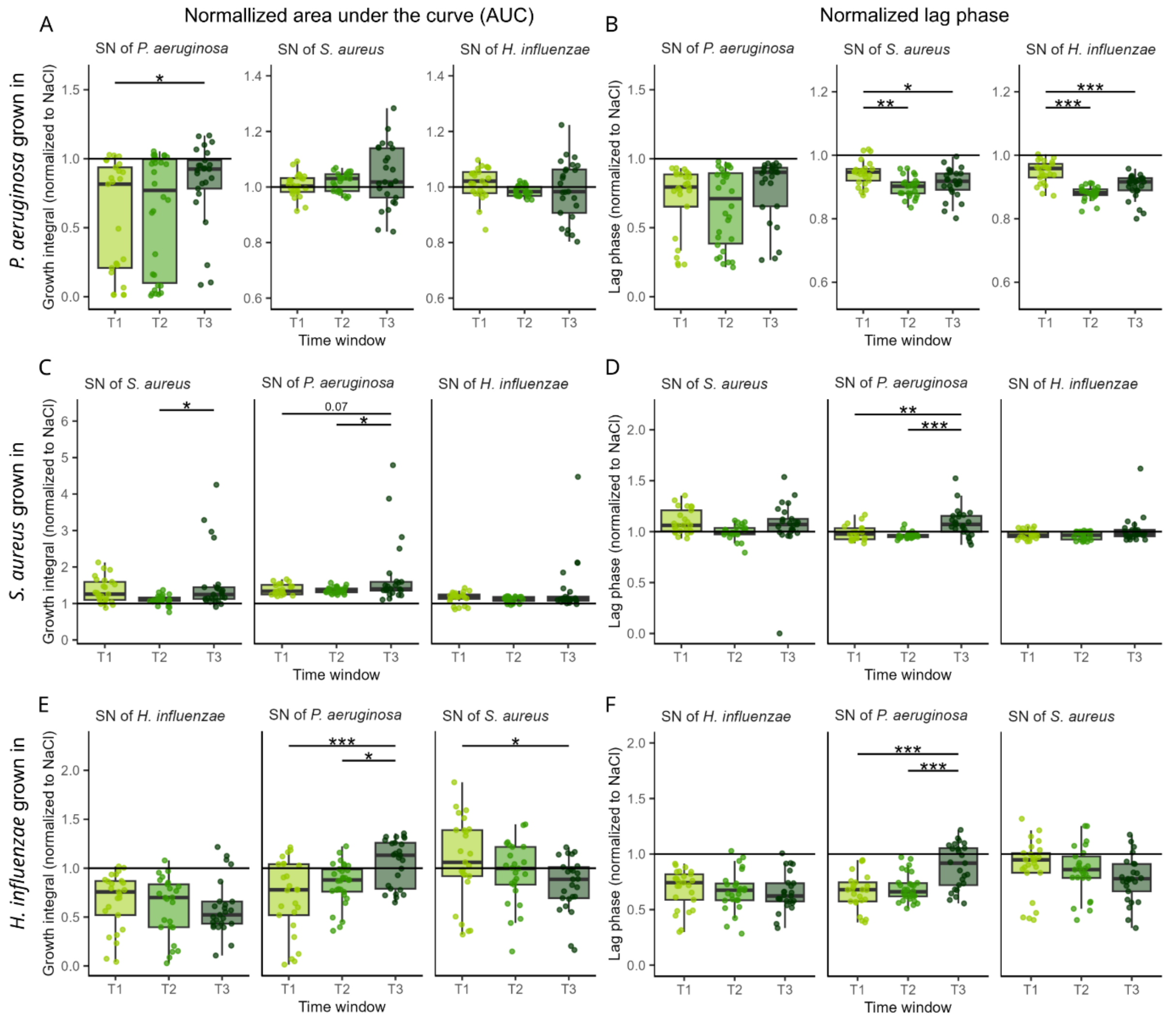


**Figure S6 | Within time point supernatant assay for studying the inhibitory potential between the clinical isolates. (A+B)** Growth integral (A) and lag phase (B) of *P. aeruginosa* in its own supernatant or the supernatant of *S. aureus* or *H. influenzae*. **(C+D)** Growth integral (C) and lag phase (D) of *S. aureus* in its own supernatant or the supernatant of *P. aeruginosa* or *H. influenzae*. **(E + F)** Growth integral (E) and lag phase (F) of *H. influenzae* in its own supernatant or the supernatant of *P. aeruginosa* or *S. aureus*. The supernatant was harvested after 18 hours growth at 37°C degrees and 170rpm. Strains were grown quadruplicates with a starting OD600 of 0.0001 in either i) 100% ssBHI ii) 62% ssBHI + 38% NaCl or iii) 62% ssBHI + 38% supernatant. We measured OD600 every two hours for 24 hours. Asterisks denote significance levels: ***P < 0.001; **P < 0.01; *P < 0.05.

**Supplemental Figure S7**

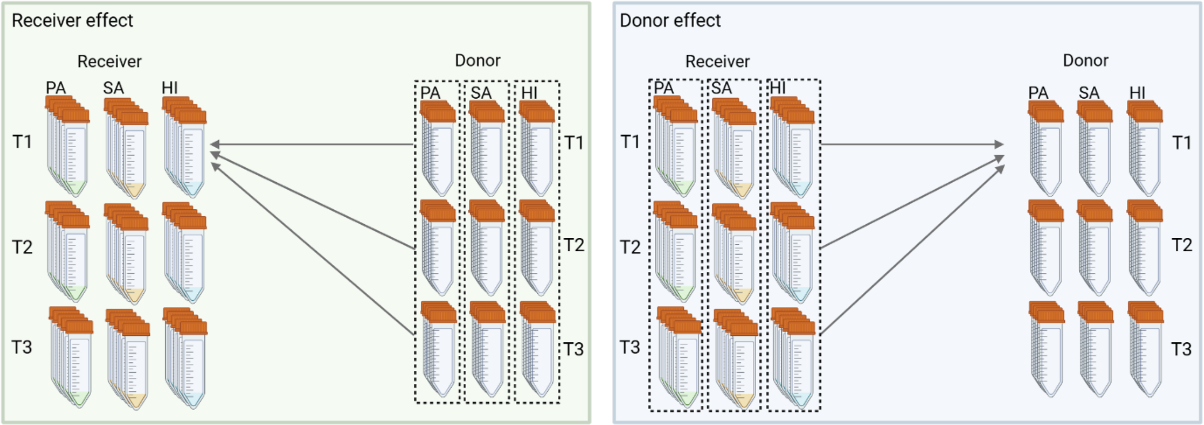


**Figure S7 | Schematic explanation of the receiver and donor effects.** Growth was measured for three receiver species: PA (*Pseudomonas aeruginosa*), SA (*Staphylococcus aureus*), and HI (*Haemophilus influenzae*) across three time windows (T1, T2 and T3). Supernatants (Donor) were also collected at T1–T3. **Receiver effect (receiver time window):** For a given receiver species, we compare growth of its T1 vs T2 vs T3 isolates while pooling across donor supernatant time window from a chosen donor species. Example: HI T1 in HI SN1/SN2/SN3 vs HI T2 in HI SN1/SN2/SN3 vs HI T3 in HI SN1/SN2/SN3. This allows us to test whether receiver time window drives growth differences, independent of the supernatant time window. **Donor effect (supernatant time window):** For a given donor species, we compare the impact of its SN1 vs SN2 vs SN3 while pooling across receiver time window of the chosen receiver species. Example: PA T1–T3 in SA SN1 vs PA T1–T3 in SA SN2 vs PA T1–T3 in SA SN3. This tests whether donor time window alters its effect on growth, independent of the receiver time window.

### Supplemental Figure S8

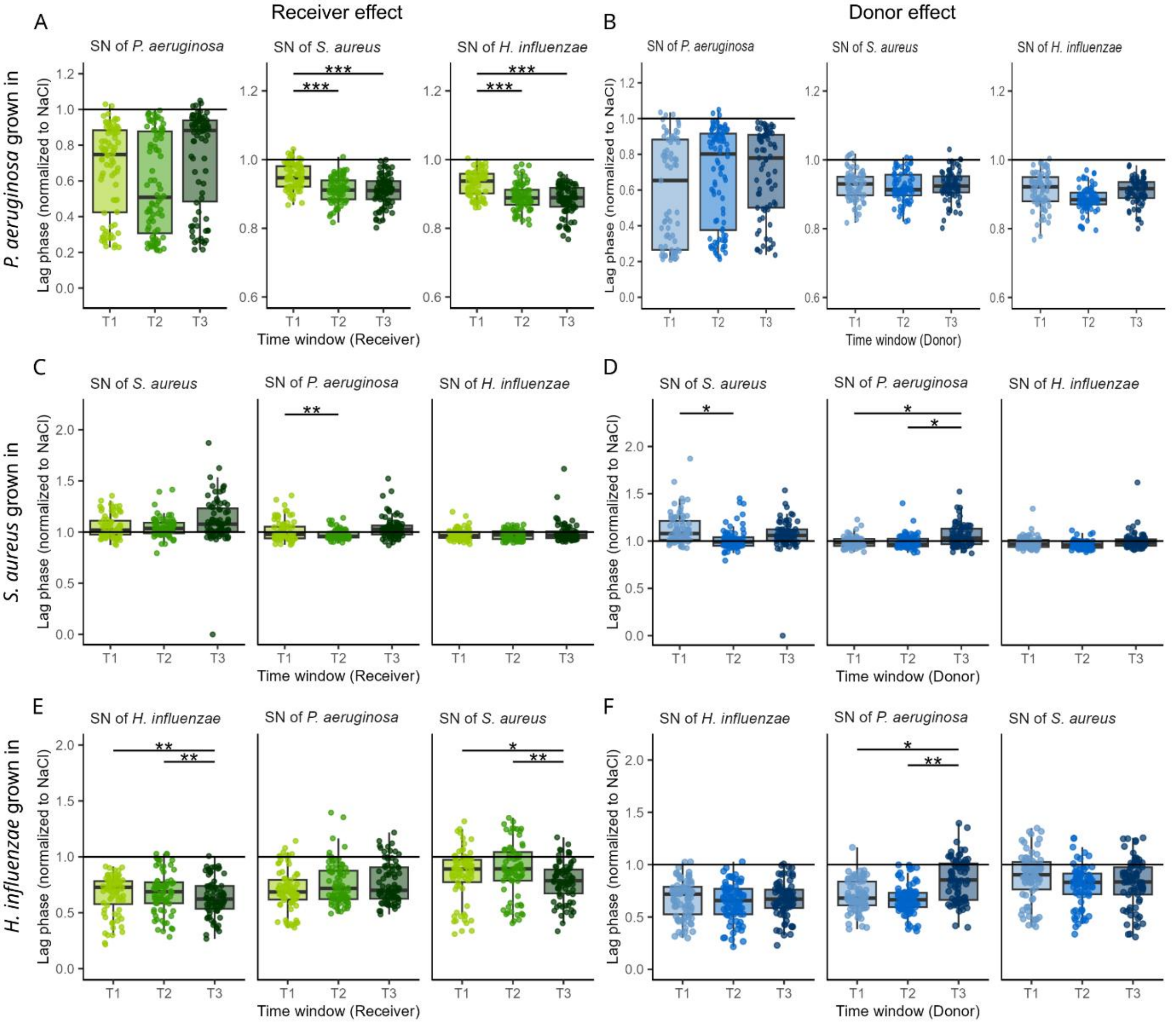


**Figure S8 | Across time window supernatant assays measuring lag phase reveal one donor effect (reduced *P. aeruginosa* supernatant toxicity at T3) and several receiver effects.** Receiver and donor effects based on normalized lag phase (lag phase relative to control condition) of isolates exposed to supernatants. The **left panels (A), (C)** and **(E)** (green boxplots) shows receiver effects: isolates from time windows T1, T2, or T3 were exposed to supernatants from all three time windows. Altered lag phase of isolates within a specific time window indicates that the isolates (receivers) differ, rather than the composition of the supernatant. The **right panels (B), (D)** and **(F)** (blue boxplots) shows donor effects: isolates from all time windows combined were exposed to supernatants sampled from T1, T2, or T3. Altered lag phase within a specific supernatant time window indicates that the composition of the supernatant (donor) differs, rather than the isolates. Panels A–C show results for different pathogens grown in the supernatants: **(A)** *P. aeruginosa*, **(B)** *S. aureus*, and **(C)** *H. influenzae*.

**Supplemental Figure S9**

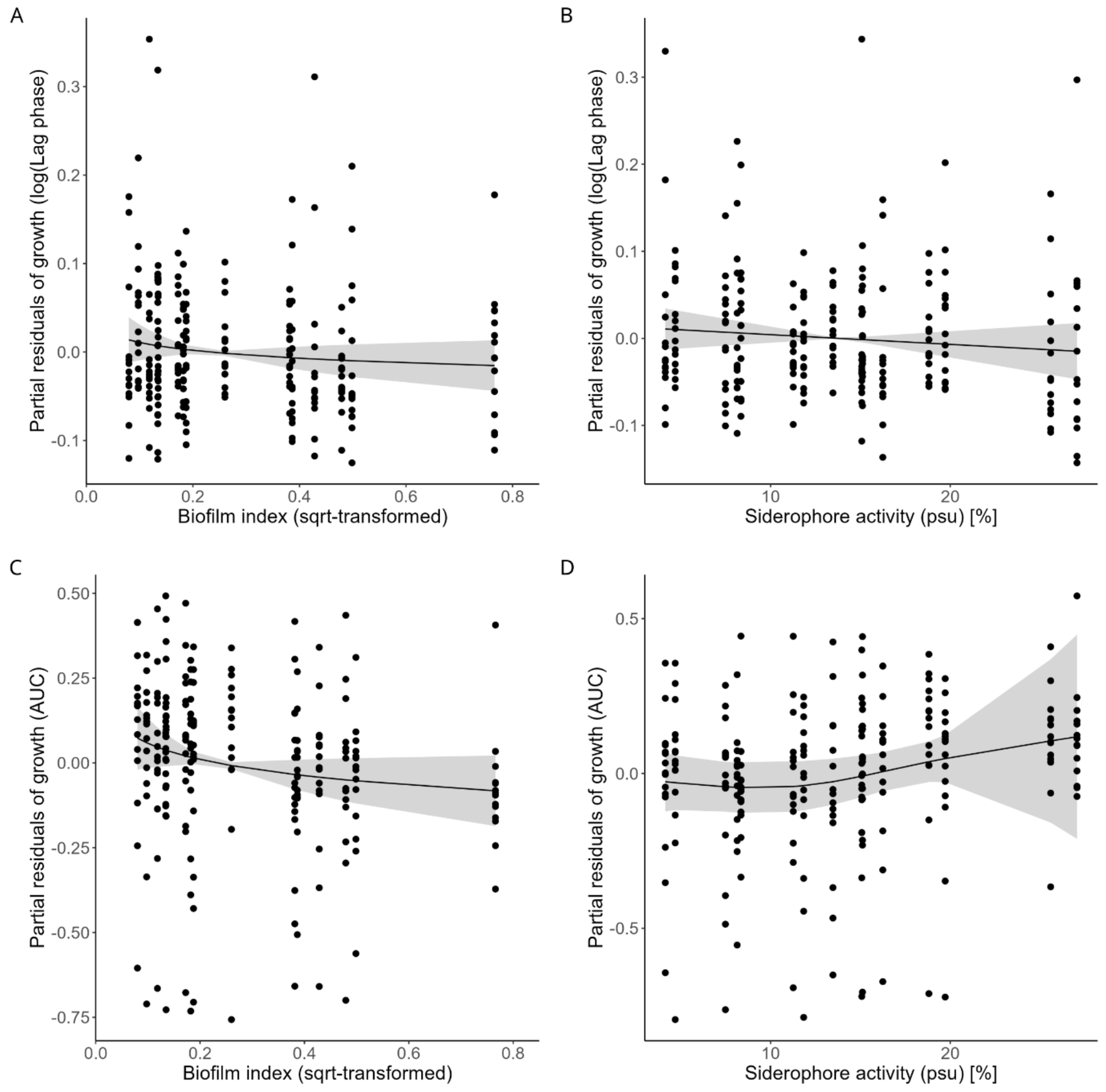


**Figures S9 | Effects of biofilm and siderophore production in *P. aeruginosa* on bacterial growth in *S. aureus* and *H. influenzae* estimated from GAMs.** Individual panels show associations between **(A)** *S. aureus* normalized lag phase (log transformed) and *P. aeruginosa* biofilm production, (**B)** *S. aureus* normalized lag phase (log transformed) and *P. aeruginosa* siderophore production, (**C)** *H. influenzae* normalized AUC and *P. aeruginosa* biofilm production, and (**D)** *H. influenzae* normalized AUC and *P. aeruginosa* siderophore production. All panels show the smooth effect of either biofilm or siderophore production with 95% confidence intervals (shaded ribbons), with points representing partial growth residuals. The GAMs included hemolysis as categorical predictor and smooth terms for log-transformed biofilm formation, siderophore and protease production.

### Supplemental Figure S10

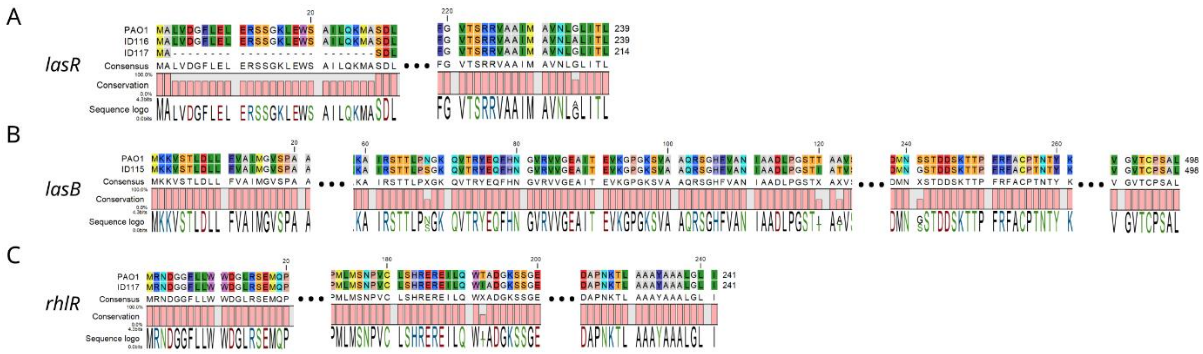


**Figure S10 | Protein sequence alignments of quorum sensing genes from *P. aeruginosa* PAO1 and clinical strains carrying SNPs. (A)** *lasR* in strains ID116 and ID117, **(B)** *lasB* gene in strain strain ID115, and **(C)** *rhlR* gene in strain ID117. All strains containing a SNP in any quorum sensing gene exhibited significantly reduced protease activity in our assay (Figure 2).

**Supplemental Table S1 | Clinical isolate used in this study**

| Species and strain ID | Person with CF ID | Time window | Child age at isolation |
|---|---|---|---|
| ***Pseudomonas aeruginosa* (PA)** | | | |
| ID100 | P14 | T1 | 1.07 |
| ID101 | P21 | T1 | 0.38 |
| ID102 | P21 | T1 | 0.59 |
| ID105 | P6 | T1 | 2 |
| ID106 | P4 | T2 | 3.15 |
| ID107 | P8 | T2 | 3.1 |
| ID108 | P6 | T2 | 4.1 |
| ID109 | P15 | T2 | 2.83 |
| ID110 | P15 | T2 | 3.01 |
| ID111 | P15 | T2 | 3.2 |
| ID112 | P6 | T2 | 3.23 |
| ID113 | P6 | T2 | 3.98 |
| ID114 | P9 | T3 | 5.49 |
| ID115 | P22 | T3 | 5.84 |
| ID116 | P22 | T3 | 6.29 |
| ID117 | P1 | T3 | 5.37 |
| ID118 | P6 | T2 | 5.03 |
| ID119 | P6 | T3 | 5.63 |
| ID120 | P6 | T3 | 7.01 |
| ***Staphylococcus aureus* (SA)** | | | |
| ID201 | P11 | T1 | 0.25 |
| ID202 | P7 | T1 | 1.4 |
| ID204 | P14 | T1 | 0.28 |
| ID205 | P14 | T1 | 1.06 |
| ID208 | P17 | T1 | 0.25 |
| ID209 | P18 | T1 | 0.25 |
| ID210 | P18 | T1 | 0.38 |
| ID211 | P17 | T1 | 0.9 |
| ID212 | P20 | T1 | 0.28 |
| ID213 | P19 | T1 | 0.16 |

| | | | |
|---|---|---|---|
| ID214 | P19 | T1 | 1.16 |
| ID215 | P21 | T1 | 0.46 |
| ID219 | P6 | T2 | 3.14 |
| ID220 | P23 | T2 | 3.42 |
| ID221 | P3 | T2 | 3.66 |
| ID222 | P5 | T2 | 3.43 |
| ID223 | P6 | T2 | 3.88 |
| ID224 | P5 | T2 | 3.98 |
| ID225 | P3 | T2 | 4.49 |
| ID226 | P22 | T2 | 3.25 |
| ID227 | P11 | T2 | 3.7 |
| ID228 | P12 | T2 | 3.11 |
| ID230 | P9 | T2 | 4.54 |
| ID231 | P12 | T2 | 4.39 |
| ID232 | P14 | T2 | 3.99 |
| ID233 | P14 | T2 | 4.64 |
| ID235 | P3 | T3 | 6.35 |
| ID236 | P5 | T3 | 6.14 |
| ID237 | P22 | T3 | 6.08 |
| ID238 | P22 | T3 | 6.17 |
| ID239 | P9 | T3 | 6.26 |
| ID240 | P2 | T3 | 5.21 |
| ID241 | P3 | T3 | 7.17 |
| ID242 | P23 | T3 | 7 |
| ID243 | P6 | T3 | 7.16 |
| ID244 | P23 | T3 | 7.08 |
| ID246 | P6 | T3 | 7.82 |
| ID247 | P9 | T3 | 7.77 |
| ID248 | P9 | T1 | 0.76 |
| ID249 | P12 | T3 | 6.12 |
| ID250 | P22 | T2 | 4.58 |
| ID251 | P16 | T1 | 0.17 |
| ID252 | P2 | T1 | 0.72 |
| ID62 | P6 | T1 | 1.5 |

***Haemophilus influenzae* (HI)**

| | | | |
|---|---|---|---|
| ID1 | P1 | T1 | 1.1 |
| ID3 | P1 | T3 | 6.6 |
| ID8 | P2 | T3 | 6.67 |
| ID10 | P3 | T2 | 4.58 |
| ID11 | P3 | T3 | 6.5 |
| ID21 | P5 | T2 | 3.92 |
| ID22 | P5 | T3 | 6.58 |
| ID33 | P7 | T3 | 5.83 |
| ID35 | P8 | T1 | 1.67 |
| ID36 | P8 | T2 | 2.92 |
| ID37 | P8 | T3 | 5.83 |
| ID41 | P10 | T2 | 4.42 |
| ID42 | P10 | T3 | 6.61 |
| ID45 | P11 | T2 | 4.34 |
| ID46 | P12 | T2 | 3.3 |
| ID47 | P12 | T3 | 5.81 |
| ID55 | P21 | T1 | 0.27 |
| ID58 | P23 | T2 | 4.43 |
| ID60 | P23 | T3 | 7.22 |
| ID61 | P13 | T1 | 1.3 |
| ID63 | P6 | T1 | 2.1 |

**Supplemental Table S2 | Antibiotic concentrations used for MIC assays.** The range of antibiotic concentrations (µg/mL) for each species used in this study.

| Antibiotic | *P. aeruginosa* | *S. aureus* | *H. influenzae* |
|---|---|---|---|
| Ampicillin | 4000, 2000, 1000, 500, 250, 125, 62.5, 31.25, 15.625, 7.8125, 0 | 1, 0.9, 0.8, 0.7, 0.6, 0.5, 0.4, 0.3, 0.2, 0.1, 0 | 12, 8, 7, 6, 5, 4, 3, 2, 1, 0.5, 0 |
| Ciprofloxacin | 2, 1, 0.8, 0.7, 0.6, 0.5, 0.4, 0.3, 0.2, 0.1, 0 | 2, 1.6, 1.4, 1.2, 1, 0.8, 0.6, 0.4, 0.2, 0.1, 0 | 0.0312, 0.0156, 0.0078, 0.0039, 0.002, 0.001, 0.0005, 0.0002, 0.0001, 6.10352E-05, 0 |
| Colistin | 15, 14, 13, 12, 11, 10, 9, 8, 7, 6, 0 | NA | 2, 1.9, 1.8, 1.7, 1.6, 1.5, 1.4, 1.3, 1.2, 1.1, 0 |
| Meropenem | 4, 3.5, 3, 2.5, 2, 1.5, 1, 0.5, 0.1, 0.06, 0 | 0.25, 0.125, 0.0625, 0.0312, 0.0156, 0.0078, 0.0039, 0.002, 0.001, 0.0005, 0 | 0.75, 0.375, 0.1875, 0.0938, 0.0469, 0.0234, 0.0117, 0.0059, 0.0029, 0.0015, 0 |
| Piperacillin–tazobactam (2:1) | 32, 16, 8, 7, 6, 5, 4, 3, 2, 1, 0 | 1.4, 1.2, 1, 0.8, 0.6, 0.4, 0.2, 0.1, 0.05, 0.025, 0 | 0.125, 0.0625, 0.0312, 0.0156, 0.0078, 0.0039, 0.002, 0.001, 0.0005, 0.0002, 0 |
| Tobramycin | 10, 9, 8, 7, 6, 5, 4, 3, 2, 1, 0 | 6, 5.5, 5, 4.5, 4, 3.5, 3, 2.5, 2, 1, 0 | 2.5, 1.8, 1.6, 1.4, 1.2, 1, 0.9, 0.8, 0.4, 0.2, 0 |

**Supplemental Table S3 | Similarity of quorum sensing gene alignments between clinical isolates and *P. aeruginosa* PAO1.** The four quorum sensing genes (*lasR*, *lasB*, *lasI*, and *rhlR*) from clinical isolates were aligned to PAO1. Values indicate sequence similarity, with 1 corresponding to 100% identity and 0.9 to 90% identity.

| Isolate ID | pwCF ID | Serogroup | Quorum sensing *lasR* | *lasB* | *lasI* | *rhlR* |
|---|---|---|---|---|---|---|
| ID106 | P4 | O6 | 1 | 1 | 1 | 1 |
| ID100 | P13 | O6 | 1 | 1 | 1 | 1 |
| ID102 | P21 | O6 | 1 | 1 | 1 | 1 |
| ID101 | P21 | O6 | 1 | 1 | 1 | 1 |
| ID117 | P1 | O6 | 0.9 | 1 | 1 | 0.99 |
| ID119 | P6 | O6 | 1 | 1 | 1 | 1 |
| ID118 | P6 | O6 | 1 | 1 | 1 | 1 |
| ID120 | P6 | O6 | 1 | 1 | 1 | 1 |
| ID112 | P5 | O6 | 1 | 1 | 1 | 1 |
| ID113 | P5 | O6 | 1 | 1 | 1 | 1 |
| ID108 | P6 | O6 | 1 | 1 | 1 | 1 |
| ID105 | P6 | O11 | 1 | 1 | 1 | 1 |
| PA7 | NA | O12 | 0.98 | 0.9 | 0.99 | 0.93 |
| ID115 | P22 | O11 | 1 | 0.98 | 1 | 1 |
| ID110 | P15 | O9 | 1 | 1 | 1 | 1 |
| ID111 | P15 | O9 | 1 | 1 | 1 | 1 |
| ID109 | P15 | O9 | 1 | 1 | 1 | 1 |
| PAO1 | NA | O5 | 1 | 1 | 1 | 1 |
| PA14 | NA | O1 | 1 | 0.98 | 1 | 1 |
| ID107 | P8 | O3 | 1 | 1 | 1 | 1 |
| ID116 | P22 | O3 | 0.99 | 1 | 1 | 1 |
| ID114 | P9 | O3 | nf | 1 | nf | 1 |

**Supplemental Table S4 | Common lab strains used in this study.** CA-MRSA: Community-acquired methicillin-resistant *S. aureus*; MSSA: Methicillin-sensitive *S. aureus*; Agr: Accessory gene regulator

| Species and strain name | Origin | Description | Reference |
|---|---|---|---|
| ***Pseudomonas aeruginosa*** **(PA)** | | | |
| PAO1 | Wound | Commonly used *Pseudomonas aeruginosa* laboratory strain. | ATCC 15692 |
| ***Staphylococcus aureus*** **(SA)** | | | |
| Cowan I | Septic arthritis | MSSA isolate. Highly invasive, but not cytotoxic. Agr-defective. | ATCC 12598 |
| 6850 | Osteomyelitis | MSSA isolate. Highly invasive, cytotoxic, and hemolytic. | ATCC 53657 |
| JE2 | Skin and soft tissue infection | USA300 CA-MRSA isolate. Highly virulent, cytotoxic, and hemolytic. | NARSA |
| ***Haemophilus influenzae*** **(HI)** | | | |
| L-378 | Lung abscess | Non-typeable strain | ATCC 49766 |
| TD-4 | Human sputum | Non-typeable strain | ATCC 49247 |
| Pittman 572 | Unknown | Serotype b (Hib), biotype I; aliases AMC 36-A-1 / / NCTC 13377 | ATCC 10211 |
